# The genomics of convergent adaptation to intertidal gravel beaches in Mediterranean clingfishes

**DOI:** 10.1101/2025.09.12.675783

**Authors:** Maximilian Wagner, Philipp Resl, Nadine Klar, Jonathan M. Huie, Iliana Bista, Shane McCarthy, Michelle Smith, Richard Durbin, Stephan Koblmüller, Hannes Svardal

## Abstract

**Introduction:** Understanding the genetic basis of widespread phenotypic convergence, particularly for complex morphological traits, remains a major challenge in evolutionary biology. The Mediterranean gravel beach clingfishes of the genus *Gouania* provide an excellent system to study this phenomenon. Within this genus, two distinct morphotypes, “slender” and “stout”, have repeatedly evolved, adapting to different microhabitats. These morphotypes differ in multiple complex traits, including body elongation, head compression, vertebral number, eye size, and the structure of the adhesive disc.

**Materials and Methods:** In a first step, to scrutinize phylogenetic convergence, we combined 3D morphometrics of the pelvic girdle and skull, with molecular species delimitation based on >660 DNA barcodes, and a phylogenomic framework based on more than 3,400 single-copy orthologs. Secondly, by employing whole-genome resequencing and a novel “convergence score” statistic, we examined genomic convergence across multiple levels: nucleotides, sequences, genes, and functional pathways.

**Results and Discussion:** While we found no evidence of large-scale genomic or protein-level convergence, we identified promising candidate regions at the level of single variants, genes, and biological pathways. Notably, a longer shared (but interrupted) haplotype around the morphogene *adam12* was associated with convergent traits. The lack of simple genomic patterns may reflect the radiation’s age and the complex genetic basis of the underlying morphological traits (e.g., eye-size, neurocranium shape). Altogether, our findings highlight the importance of assessing genomic convergence at multiple molecular levels to uncover diagnostic signals across varying evolutionary processes and timescales.

**GRAPHICAL ABSTRACT:** 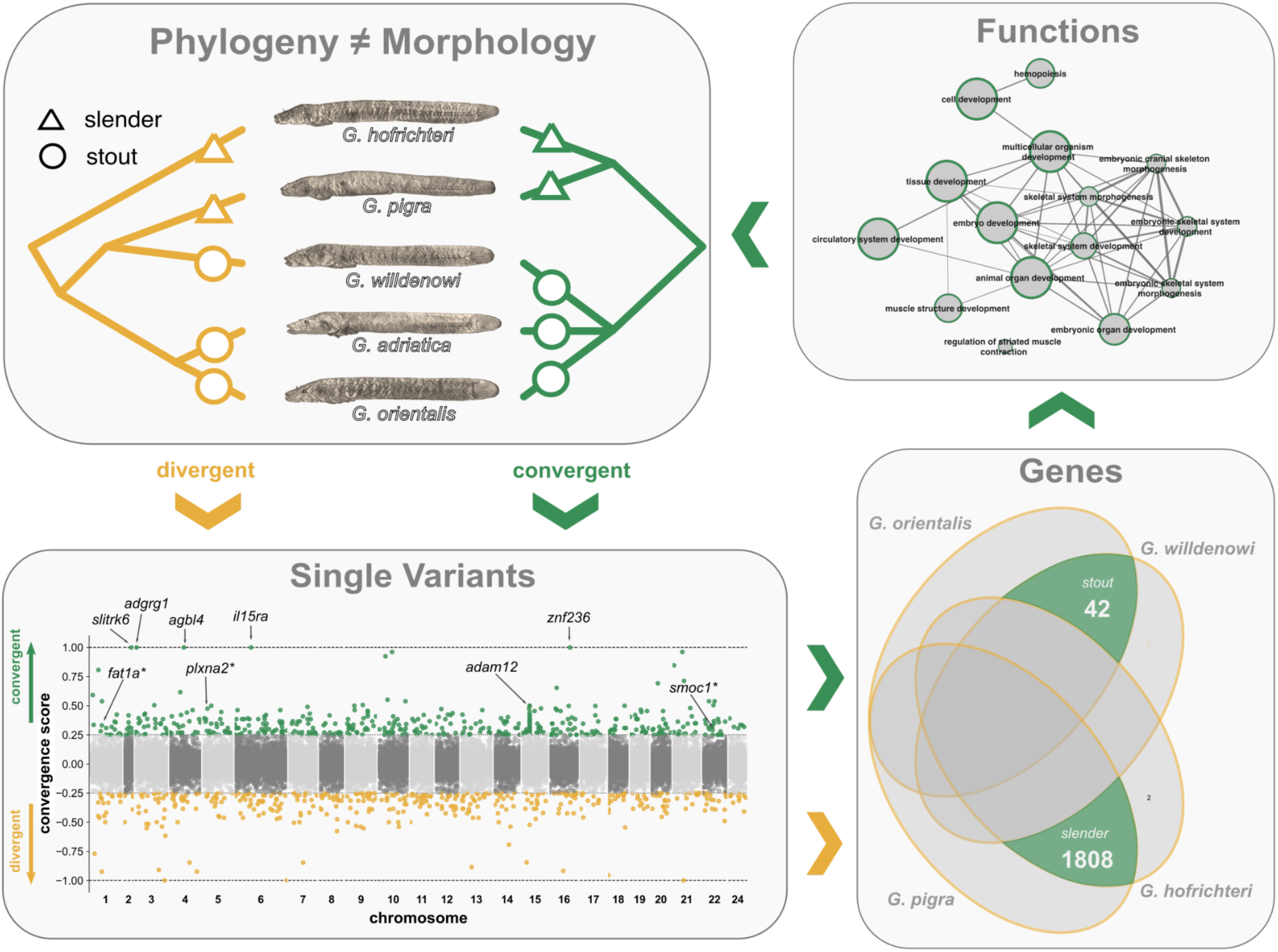

## INTRODUCTION

The independent evolution of similar phenotypic adaptations in non-sister lineages (convergent or parallel evolution) is one of the most intriguing biological phenomena and can help provide answers to fundamental questions such as the contingency of evolution and the molecular underpinnings of adaptation. Teleost fishes hold some of the most compelling examples of convergent evolution in the animal kingdom, for example in the context of marine–freshwater transitions (Thacker and Gkenas 2019; Magalhaes et al. 2021; Friedman et al. 2022), or with respect to adaptations to extreme environments (Wang and Guo 2019) such as caves (Powers et al. 2020), sulfuric springs (Tobler et al. 2011), high altitudes (Yang et al. 2021), or the deep sea (Shen et al. 2019). The accumulation of cases of convergent evolution in the context of adaptation to unstable environments is not a coincidence, but likely reflects the strength of selection and restrictedness of evolutionary outcomes under such extreme environmental conditions (Losos 2011; Xu et al. 2020).

Among the most demanding environments on the planet are the interstitial spaces of marine gravel beaches. The prevailing dynamic and hostile conditions, including tidal fluctuations, salinity changes, and especially intense mechanical forces, allow only highly adapted species to survive. Among the few vertebrates that have evolved to thrive in extreme interstitial gravel ecosystems are Japanese wormgobies (*Luciogobius*, Gobiidae) and the blunt-snouted clingfishes (*Gouania*, Gobiesocidae) endemic to the Mediterranean (Yamada et al. 2009; Wagner et al. 2019). Both gobies and clingfishes independently evolved a ventral sucking disc, but in clingfishes, the disc is also supported by the pelvic girdle, potentially enhancing evolutionary modularity and adaptive potential (Huie et al. 2022).

Compared to other species in their families, both genera have independently evolved an extreme worm-like, and partly troglomorphic appearance, which is primarily driven by an increase in vertebrae numbers and a reduction of fins and eye size (Briggs 1955; Yamada et al. 2009; Wagner et al. 2019). Furthermore, Mediterranean *Gouania* and Japanese worm gobies have further diversified in these environments along an axis of body elongation presumably adapting to distinct microhabitats (Yamada et al. 2009; Wagner et al. 2023).

Specifically, within *Gouania*, the five recently discovered species have been assigned to “stout” and “slender” morphotypes, with slender species having an increased number of vertebrae, a more compressed head shape, and smaller eyes, fins and sucking discs compared to their stout counterparts (Wagner et al. 2019; Wagner et al. 2021). While there is only a single stout to intermediate type in the western Mediterranean, separate pairs of stout and slender species co-exist in the Adriatic and eastern Mediterranean, respectively (Figure 1a). Where sympatric, the two morphotypes occupy different microhabitats, with slender species more likely to be found in beach areas with smaller gravel sizes, compared to their stout congenerics, consistent with divergent adaptation to distinct interstitial space profiles (Wagner et al. 2023). A previous multi-gene phylogeny (Figure 1b) suggested that the different morphotypes evolved repeatedly in different *Gouania* lineages (*6*). However, evidence based on the analysis of whole-genome data is lacking.

**Figure 1.**
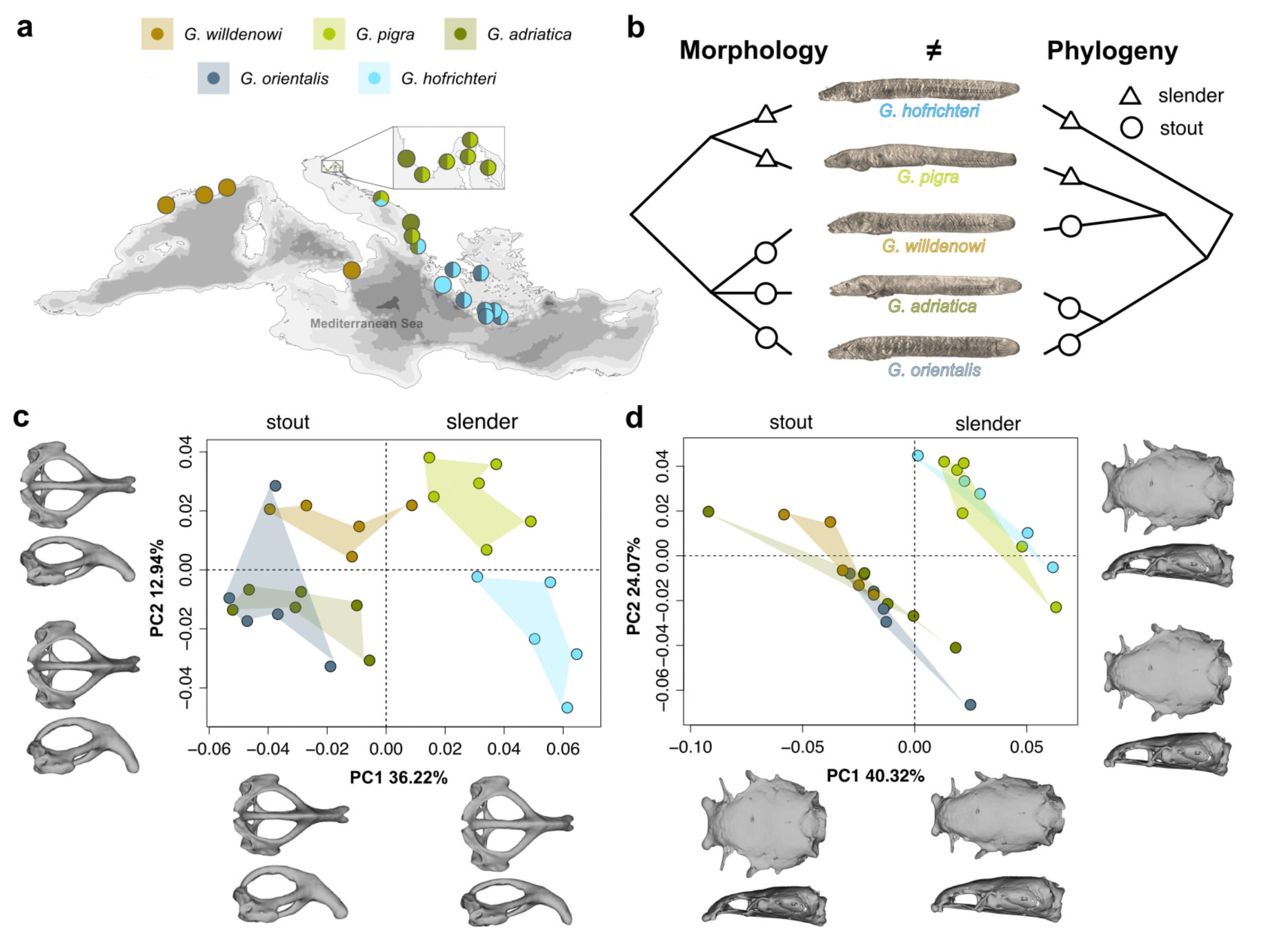
Morphological convergence, distribution and phylogenetic relationships of the blunt-snouted clingfishes (*Gouania*). **(a)** Currently known distribution ranges and sampling sites for diversity screening (see Supplementary Results/Discussion). **(b)** Previous research (Wagner et al. 2019) suggested that *Gouania* morphotypes, slender and stout evolved convergently. **(c-d)** Three-dimensional geometric morphometric analysis of the **(c)** pelvic girdle and the **(d)** neurocranium clearly separate two morphotypes, slender and stout, along the first two principal components but generally not species within morphotypes.

Convergent evolution at the molecular level is a complex question and can unfold in various ways. First, since the genetic basis of complex phenotypes is most often polygenic, some underlying loci might be shared but not others (Cerca 2023). Second, qualitatively different levels of sharing of a molecular basis can be distinguished. At the lowest level, adaptive alleles can be shared between convergently evolving lineages because they appeared once in a shared common ancestor. Such shared mutations from a single origin — referred to as “identity by descent” (IBD) in population genetics or “collateral evolution” sensu (Stern 2013) — may be the result from introgression between lineages undergoing convergent evolution, or ancestral polymorphisms that were maintained over time and later independently selected in diverging species pairs. As an alternative to IBD, mutations could appear independently in different convergently evolving lineages, either affecting the same nucleotide positions, the same genes but different nucleotides, or different genes within the same or different molecular pathways (Elmer and Meyer 2011). Distinguishing between these different alternatives is challenging (Stern 2013; Alaei Kakhki et al. 2023), but critical for our understanding of the processes behind evolutionary convergence.

In this study, we combine 3D-morphometrics, molecular species delimitation based DNA barcodes and genomics to scrutinize convergent evolution in *Gouania* and investigate its molecular underpinnings. We specifically focus on the different levels of molecular convergence, testing for signatures at the level of nucleotides, sequences, genes and pathways.

## RESULTS/DISCUSSION

### Parallel phenotypic divergence among *Gouania* lineages confirmed by 3D geometric morphometrics and phylogenomics

A number of different terms have been employed to describe the evolution of similar phenotypic adaptations in non-sister lineages (Arendt and Reznick 2008; Stern 2013; Alaei Kakhki et al. 2023; Cerca 2023). The most prominent – convergent and parallel evolution – have variably been used to distinguish between different phylogenetic levels, molecular bases, and trajectories of phenotypic change (Arendt and Reznick 2008; Elmer and Meyer 2011; Stern 2013; Alaei Kakhki et al. 2023; Cerca 2023). In this study, we will not distinguish between the terms (following (Arendt and Reznick 2008)), but focus on questions about the (shared) molecular basis of phenotypic convergence.

To confirm the previously suggested phenotypic dichotomy of slender and stout morphotypes of *Gouania* (Wagner et al. 2019), we investigated 26 linear measurements (Figure S1), as well as three-dimensional shape differences of the pelvic girdle and the neurocranium (Figure 1c,d) in a sample of 27 individuals of the five recognized species. Both the 3D geometric morphometric (GEMO) and the linear analyses clearly separated the five species into the two main morphotypes, slender and stout (Figure S1; Figure 1c,d). Overall, the first two PCs explained 49.16% and 64.39% of shape variation for the pelvic girdle and the neurocranium, respectively. Shape differences of the pelvic girdle reflected changes in the width of the girdle with slender types showing an overall wider pelvis compared to stout morphotypes (Figure 1c). Notably, *G. willdenowi*, the stout species not co-occurring in sympatry with a slender species, occupied a slightly more intermediate area of the morphospace (Figure 1c). For the neurocranium, the first principal component mainly loaded on differences representing a narrower dorsolateral, but a wider dorsoventral shape in slender compared to stout morphotypes (Figure 1d). Finally, since the adhesion capacity of clingfish correlates with the area or size of the sucking disc (Huie et al. 2022), we also compared disc sizes among species and found them to be significantly smaller in slender morphs (Figure S2).

Our morphological results nicely align with previous findings that show a more compact, but elongated body shape is preferred by slender *Gouania* inhabiting smaller interstitial spaces (Wagner et al. 2019; Wagner et al. 2021; Wagner et al. 2023). Respectively, the overall smaller but wider pelvic girdle in slender *Gouania* species might indicate adaptations to narrower interstitial spaces, where suction ability becomes less important due to smaller grain sizes compared to those occupied by stout *Gouania* species (Wagner et al. 2023).

The interpretation of morphological convergence in the genus might be flawed by overlooked (cryptic) diversity. To address this, we analysed mitochondrial cytochrome-c-oxidase I (COI) DNA barcodes from 668 individuals of 23 populations of all described species across previously recorded sites (Figure 1a). The results strongly support the validity of the applied five species taxonomic model for *Gouania* (Figure S3; see Supplementary Results/Discussion).

Previous phylogenetic analysis based on nine nuclear loci and one mitochondrial marker suggested that slender–stout divergence happened at least two times in parallel in *Gouania* (Wagner et al. 2019). To scrutinize this result, we assembled draft genomes for four *Gouania* species and the shore clingfish (*Lepadogaster lepadogaster*), the presumed sister-lineage to *Gouania* (Conway et al. 2020) and used those genomes together with recent reference genomes of *Gouania adriatica* and the jewelled blenny, S*alaria fasciatus* (Rhie et al. 2021) in the phylogenomic analysis, based on 3,406 single-copy orthologous BUSCO genes (Table S1, S2).

Both a concatenated maximum likelihood (ML) phylogeny (Figure 2a) and an ASTRAL species tree composed from single gene trees consensus of single gene trees (Figure S4) suggested that the slender *G. hofrichteri* is the sister group of the Adriatic and Eastern Mediterranean stout species, *G. adriatica* and *G. orientalis*, while the Western Mediterranean stout *G. willdenowi* and the Adriatic slender *G. pigra* form a sister group basal to those lineages. However, despite obtaining high bootstrap support (and quartet scores from the ASTRAL tree) for all nodes, we observed low site- and gene-wide concordance factor values, especially in deeper nodes close to the root (Figure 2a). Furthermore, a neighbor-joining tree of pairwise differences showed different phylogenetic relationships, placing *G. hofrichteri* as the global outgroup to all other species (Figure S5; Figure 2b). Overall similar topological incongruences were observed in a previous study based on a concatenated tree using nine nuclear loci and one mitochondrial marker, which, only after inclusion of the mitochondrial marker, resolved *G. hofrichteri* as sister to all other *Gouania* species (Wagner et al. 2019).

**Figure 2.**
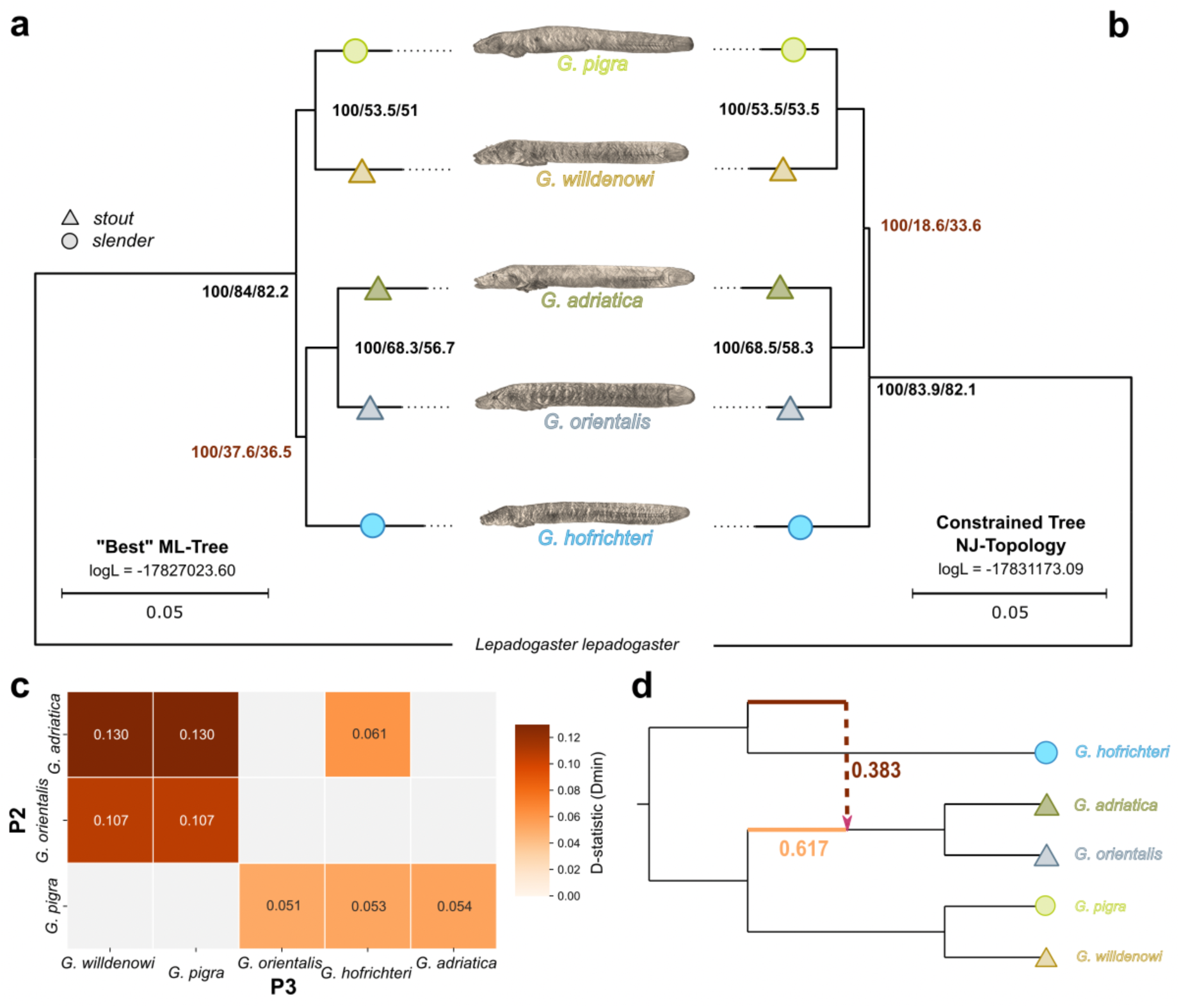
Phylogenetic incongruence at the base are most likely explained by ancient gene-flow. **(a)** The “best” Maximum-Likelihood (ML) tree estimation from a concatenated dataset of 3,406 single-copy BUSCO genes. **(b)** ML-based constrained tree to the Neighbour-Joining topology (Figure S5), revealed a slightly different topology with *G. hofrichteri* being the global outgroup to all other *Gouania* species, albeit with lower overall log-Likelihood values. For each node, the bootstrap support values, the gene- and site-concordance factors (i.e., the fraction of genes or single nucleotides that support a certain node) are given. The trees are further annotated according to slender (circles) and stout (triangles) morphotypes. **(c)** D-statistics (Dmin) show an excess allele sharing between different *Gouania* lineages (all values are significant after correction for multiple testing - Table S3). **(d)** The best-scoring phylogenetic network (restricted to one hybrid edge) calculated with PhyloNetworks based on all maximum-likelihood trees inferred from 3406 BUSCO-loci. The network shows a 38.3% introgression from the ancestor of *G. hofrichteri* into the ancestor of *G. adriatica* and *G. orientalis*.

Since low concordance factors and inconsistencies between tree building methods can be a sign of complex (network-like) phylogenetic relationships, we next investigated whether gene flow contributed to the evolution of *Gouania*. Indeed, ABBA-BABA tests of excess allele sharing revealed strong signals of network-like evolution, consistent with *G. hofrichteri* constituting an outgroup lineage to other *Gouania* species that had genetic exchange with an ancestor of *G. adriatica* and *G. orientalis* (Figure 2c; Table S3). The directionality of such a gene flow event cannot be distinguished in ABBA-BABA tests, but a reconstruction based on phylonetworks confirmed a contribution from a lineage related to the ancestor of *G. hofrichteri* into the ancestor of *G. adriatica* and *G. orientalis*. We thus consider this the most likely evolutionary scenario (Figure 2d). Importantly, all phylogenetic inference and gene flow results as well as phylogenetic hypothesis testing support at least two slender–stout transitions within *Gouania* and thus the presence of convergent evolution (Figure S6; Table S3; Table S4; see Supplementary Results/Discussion).

### No evidence for accelerated protein evolution

To investigate the genomic basis of phenotypic convergence, we analysed population level whole-genome sequencing data for 58 specimens (10 to 13 individuals per species). Our final variant callset included 22 million biallelic variants that passed filtering criteria (19 million SNPs, 3 million indels) with most variation observed in non-coding parts of the genome such as introns, and intergenic regions (Figure S7). To test whether ecomorph divergence was accompanied by accelerated protein evolution, we computed the genome-wide neutrality index for all species comparisons. Out of six slender–stout comparisons, two showed a neutrality index significantly smaller than 1.0 consistent with a genome-wide excess of positive selection. However, the neutrality indices for slender–stout comparisons were not significantly lower compared to within ecotype comparisons (Figure S8; NI_TG_, *p =* 0.7542). Consistent with this, only a maximum 23 out of 22,687 genes showed significant evidence for positive selection having acted on them, with the largest numbers consistently found in comparisons that involved *G. hofrichteri* (Figure S9).

### No signals of convergent evolution at large genomics scales

To test whether larger genomic regions show signals of convergent evolution between slender– stout species pairs, we investigated summary statistics in genomic windows of 20,000 base pairs (bp). The cluster separation score (CSS), a measure of convergent evolution (Jones et al. 2012; Miller et al. 2019), did not show any significant outliers. This suggests the absence of longer shared haplotypes between convergently evolving species. Presence of longer shared haplotypes would have been expected if convergence were due to relatively recent genetic introgression.

Even without a recent shared origin, the same genomic regions could have responded to selection in the different slender–stout species pairs independently, thus leading to shared signals of genetic divergence across diverging ecomorphs. To test for this, we calculated net nucleotide divergence (*D_a_*) between slender–stout species pairs (Figure S10) and investigated the overlap of signals of high divergence across comparisons. We got 1,998 and 2,058 outlier windows for the two pairwise comparisons, respectively. In total 145 outlier windows, corresponding to 294 genes, were shared between the two comparisons, which is similar to what would be expected simply by chance (i.e., 117 windows at *p* > 0.05; Figure S10), suggesting that genomic signals of slender–stout divergence are not shared at larger genomic scales.

### Single-nucleotide changes point towards convergent adaptation in morphological and neurological developmental processes

To investigate whether single genomic variants reflected convergent evolutionary patterns, we developed the “convergence score” (Figure 3a), a statistic that measures whether allele frequency shifts at SNPs happened convergently (i.e., increased frequency of the same allele in same-morph comparisons, convergence score > 0) or divergently (increased frequency of the same allele in opposite-morph comparisons, i.e., divergence, convergence score < 0). Overall, of the 22 million variants, 1.4% showed a non-zero convergence score, corresponding to allele frequency shifts in both ecomorph comparisons. Among those, there was a small but significant excess of convergent changes (156,640 vs. 154,608, binomial test *p* = 0.000272).

**Figure 3.**
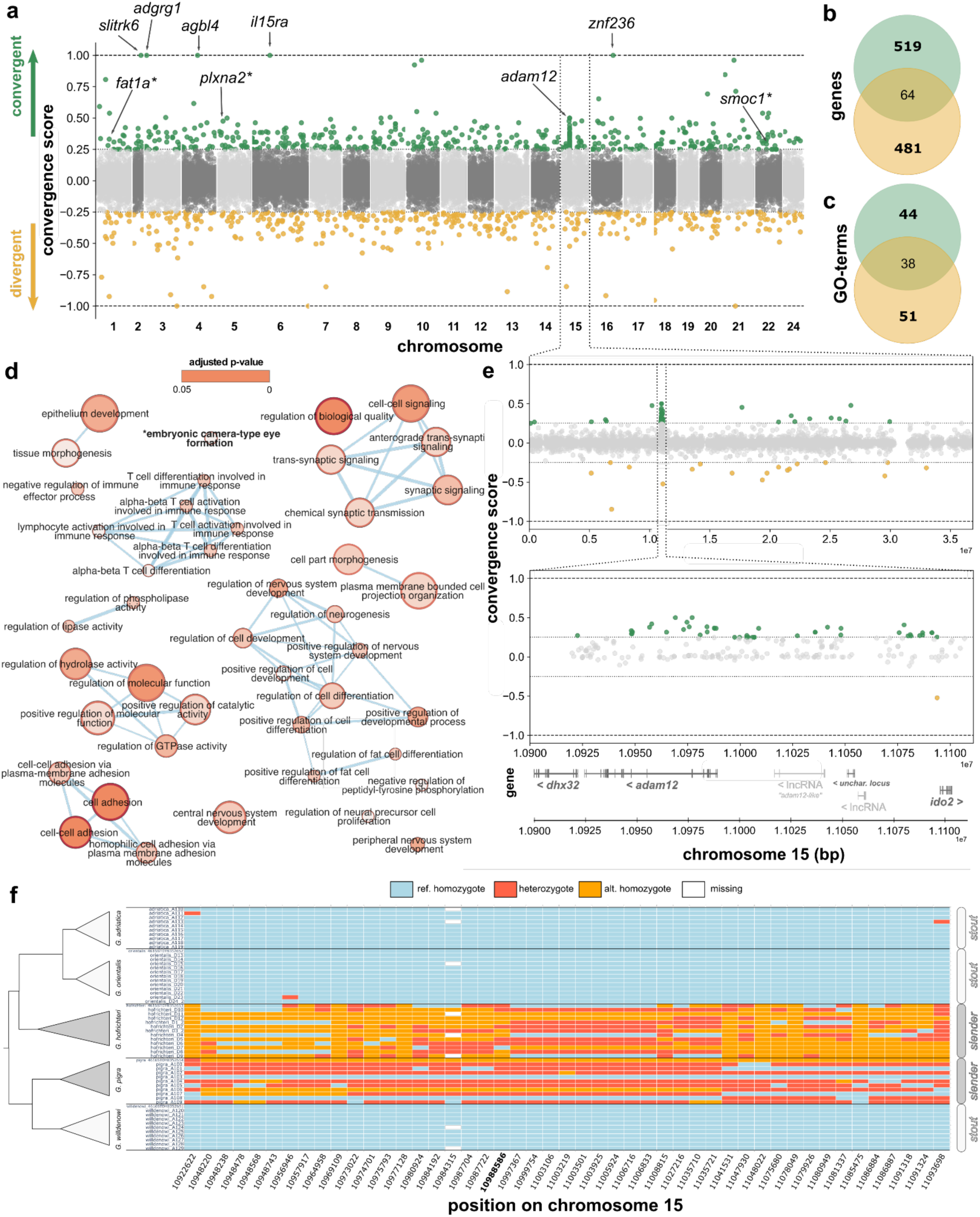
Homologous changes (snps and indels) and associated genes and functional enrichments based on convergent allelic frequency shifts (convergence score). **(a)** Convergence score values for each variant (snps and indels) along the 23 chromosomes of the *Gouania* genome. Negative values indicate divergent allele-frequency shifts whereas positive values show convergent allele frequency shifts (i.e., consistent with morph assignment). Values higher than 0.25 and lower than -0.25 are highlighted in green and yellow, respectively (see Material and Methods). Note that in the figure only the downstream names of genes are given. **(b)** Venn diagram of intersection between genes associated with outlier regions for negative and positive scores and **(c)** overlap between biological functional enrichment categories obtained for the unique 519 and 481 genes. **(d)** GO-enrichment network showing the relationships among the 44 unique GO categories associated with positive changes. **(e)** Chromosome 15 details and highlights of the *adam12* haplotype. **(f)** Genotype plot of all significant positive outliers in the *adam12* region (green dots shown in (e); convergence scores >= 0.25). The missense mutation on *adam12* exon 3 is highlighted in bold (position 10,988,596).

Among the five variants convergently fixed (convergence score = 1), there was a mutation on chromosome 2 upstream of *slitrk6*, a gene expressed early in the development of the nervous system of zebrafish and throughout adulthood, whereby expression has been mainly detected in the otic vesicle (Round et al. 2014). The gene has been found to impact retinal development and has been associated with myopia in mice and humans (Tekin et al. 2013). Convergently fixed changes also included an intron variant on chromosome 3 in the gene *adgrg1*, important in brain development in humans (Singh and Lin 2021), a variant on chromosome 4 upstream of the gene *agbl4* – a paralog of *abgl5* that is a promising candidate to cause *retinitis pigmentosa*, a sickness that could lead to the progressive loss of vision up to blindness (Kastner et al. 2015) – and an intergenic variant on chromosome 16 between *mbpb* and *znf236*. Finally, a convergently fixed variant on chromosome 6 corresponded to a premature stop codon in *il15ra*, a gene that plays a crucial role in the immune response in fishes (Jiang et al. 2024).

To identify further candidate regions, we focused on the variants with convergence scores of at least 0.25 in absolute value, corresponding to a total allele frequency shift of at least 0.5 (Figure 3a). The resulting 994 outlier sites again showed a small but significant excess of convergent changes (535 vs 459, binomial *p* = 0.01732). Associated with these outliers, we detected 519 and 481 genes that only showed positive and negative outliers, respectively (Figure 3b). For the 519 genes associated with positive convergence scores we found 44 uniquely enriched GO-terms (i.e., significantly associated with genes of positive convergence score outliers but not with negative ones; Figure 3c). These included neurological terms such as *regulation and development of central nervous system, cell- or synaptic-signalling* and *embryonic camera-type-eye formation*, but also one larger term representing *T-cell* differentiation (Figure 3d; Table S5).

The neurological enriched GO-terms identified are interesting, because *Gouania* (and potentially also other clingfishes) lack immunoglobulin genes which are essential for the adaptive immune response in other organisms (Mirete-Bachiller et al. 2021). The engagement of T-cells could offer an alternative response to immunological challenges experienced by *Gouania*. Alongside the stop-codon-inducing mutation in *il15ra*, our results may suggest selection toward similar pathogen pressures acting on both slender and stout morphs within their respective microhabitats.

Nonetheless, the overall predominance of non-coding changes among convergently fixed and convergence outlier variants suggest an important role for gene regulatory changes affecting the morphological development of slender and stout ecotypes. Supporting this, we found convergent evolution outlier genes to be strongly enriched for being associated with binding sites of transcription factors (TFs) playing important roles in development, including members of the *pax*, *hox*, *fox*, *meis* and *tcf* TF families (Table S5). For instance, *pax6* is a strong candidate for explaining eye loss in cave fishes (Strickler et al. 2001) and *hox* genes are important during vertebrate development.

Finally, the appearance of neurological and visual system related development candidate genes as well as the enrichment of associated gene ontologies (e.g., *embryonic camera-type-eye formation*) among convergence outliers is not unexpected given the significant differences in eye sizes between slender and stout morphotypes (Wagner et al. 2019; Wagner et al. 2021). Compared to stout species, slender *Gouania*, which inhabit narrower and darker interstitial spaces,encounter a lower light environment and relatively stronger mechanical forces. Consequently, convergently evolved retardations of eye development in slender species might follow a similar trajectory as eyesight loss in cave fishes compared to their surface-dwelling relatives (Jeffery 2005). Such changes might constitute an energy saving mechanism, but might also be an adaptation to reduce risk of mechanical injury, as open wounds in fishes quickly become infected by fungi or bacteria. Indeed, throughout the sampling period, we often found adult individuals that lost one or even both eyes (Figure S11).

### Extended region of convergent evolution around *adam12*

Consistent with the absence of large windows of genomic convergence seen above, genetic variants with high convergence scores generally did not cluster in specific genomic regions (Figure 3a). However, a notable exception was a peak of several convergent variants on chromosome 15, which were mainly in and downstream of the gene *adam12* (Figure 3e). This gene encodes a transmembrane protein with various functions such as cell-to-cell interactions, cell adhesion, and intracellular signalling (Kveiborg et al. 2008). Most of the variation in this region (again) corresponded to non-coding nucleotides (Figure S12) but we found a single missense mutation in the third exon of *adam12* (chromosome 15:10,988,596 or *adam12* exon 3: 43 AA position), which affected the two slender morphs (Figure S13). In this exon we detected two and one more private missense mutations for the slender *G. pigra* and *G. hofrichteri*, respectively (Figure S13). Additionally, the region also included several non-coding RNAs (LncRNA) downstream of *adam12* and a gene upstream of *adam12* (ENSGWIG00000018693; *adam12*-like) that was annotated as lncRNA but showed high sequence similarity to *adam12* (Figure 3e) consistent with a relatively complex evolutionary history of these genomic regions that involves gene duplication. However, we found no substantial difference in relative sequencing coverage among species in this region, suggesting that the observed signal is not due to copy number variation (Figure S14).

The striking accumulation of convergently evolved genetic variants (convergence score >=0.25) in the *adam12* region makes it unlikely that convergent changes evolved independently among species pairs, but rather indicates common ancestry of the mutations present in species of the same morphotype (Figure 3f). However, the fact that haplotypes are overall still relatively divergent (Figure S15) speaks against recent transfer through introgression, but rather suggests a relatively old shared origin, potentially accompanied by some form of recombination suppression that helped to link variants together.

In zebrafish, *adam12* plays a role in juvenile growth, particularly by controlling genes affecting bone and cartilage development. *Adam12*-deficient fish exhibit reduced body growth without major developmental defects (Tokumasu et al. 2016). In cichlid fishes, this gene is linked to soft tissue development involved in hypertrophy and craniofacial development, including parts of the neurocranium (Conith et al. 2018; Conith et al. 2024). Furthermore, *adam12*-deficient zebrafish exhibit smaller ligaments compared to wild types and show truncated coronoid processes of the mandible (Conith et al. 2018). Overall, this makes *adam12* a strong candidate gene for convergent adaptation of *Gouania* morphotypes.

### Non-homologous fixed changes associated with developmental processes

The convergence score investigated above does not capture non-homologous molecular convergence, that is, changes leading to convergent phenotypic adaptation that affect different parts of the same gene or different genes in the same pathway (Elmer and Meyer 2011). To investigate the potential presence of such changes, we identified genes with independently fixed genetic variants (i.e., sites corresponding to a fixed difference between one species and all others) and tested for overlap within morphotypes in terms of the gene content and function (Figure 4). We focused on one pair each of convergently evolved slender and stout species (i.e., excluding *G. adriatica* from the analysis), but an analysis including all species yielded similar results (Figure S16; see Supplementary Results/Discussion).

**Figure 4.**
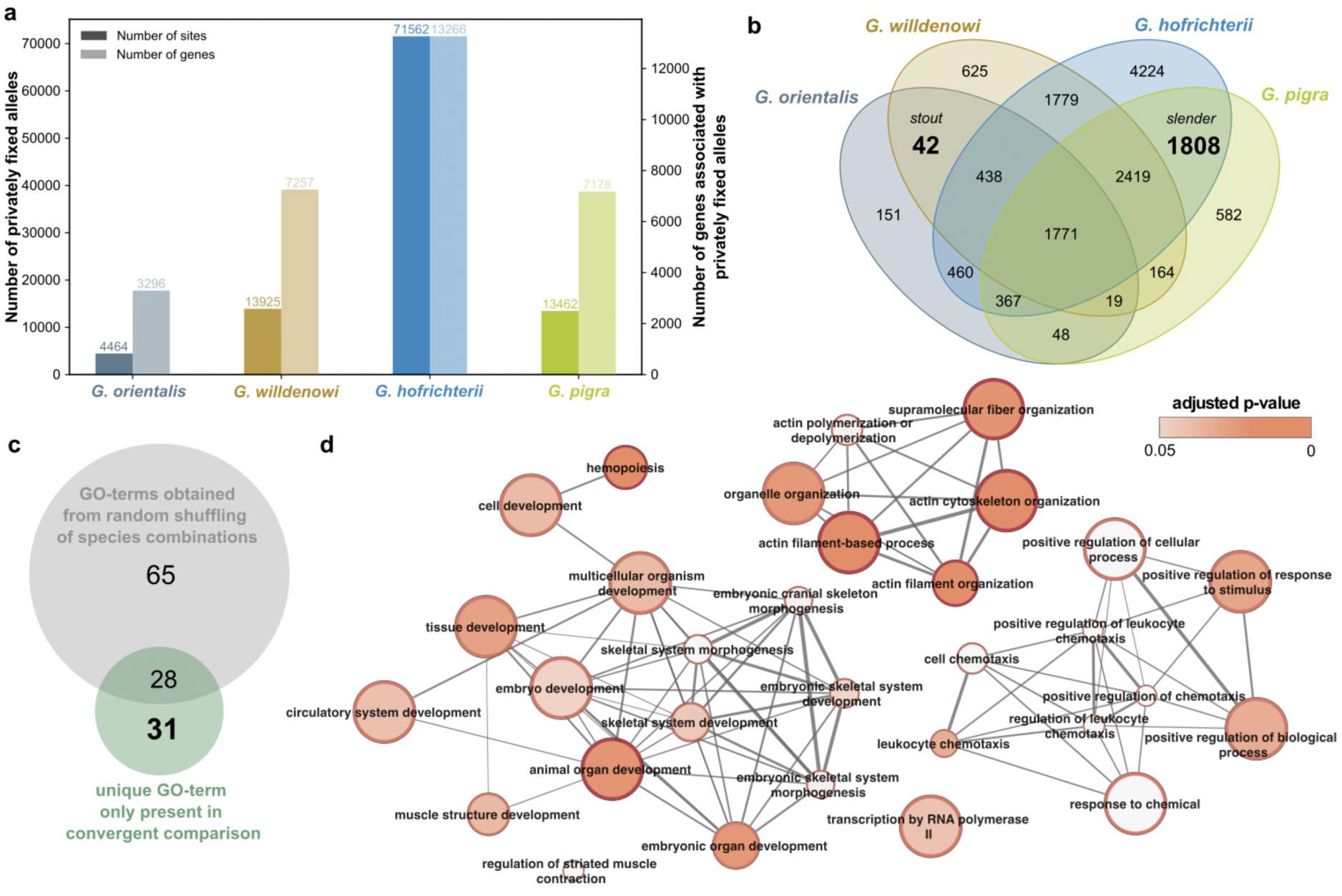
Non-homologous fixed variation involved in convergent evolution of the genus *Gouania*. **(a)** Number of sites where one allele is fixed in the given species but absent in all others, as well as the associated number of genes. **(b)** Venn diagram of number of genes with differentially fixed mutations. Highlighted are the 42 and 1,850 genes associated with differentially fixed mutations that overlap between the stout (*G. willdenowi*, and *G. orientalis*) and slender (*G. hofrichteri* and *G. pigra*) morphotypes, respectively. **(c)** After conducting a GO enrichment analysis of random comparisons of gene sets of morphotypes only 31 unique GO terms remain for convergent comparisons. **(d)** Network showing the 31 GO-terms unique to convergent patterns.

The number of differentially fixed variants and corresponding genes (“differential fixation candidate genes”) were substantially different between species (Figure 4a), with *G. hofrichteri* showing by far the largest number of fixed differences consistent with its inferred basal phylogenetic position (Figures 2a, 4a). We found that 42 genes showed fixed differences in both stout morphotypes but in neither of the slender morphs, while 1,808 genes showed the opposite pattern (Figure 4b). The larger number in the latter category can be explained by the overall larger number of genes identified in *G. hofrichteri*. Overall, there was no significant over-representation of genes shared between within-morphotype comparisons compared to between morphotype comparison (permutation; *p* = 0.33).

For the total 1,850 morphotype-specific differential fixation candidate genes (1,808 slender + 42 stout), we detected 59 significantly enriched biological process categories, 31 of which were not found in any permutation of slender and stout labels (see Methods; Figure 4c). Several of these unique GO terms involved embryonic organ and skeletal development (e.g., *skeletal system development, embryonic cranial skeleton morphogenesis*) (Figure 4d; Table S6). Together with the fact that the identified genes again showed significant associations with transcription factor binding motives (Table S6), further supports broad convergence in regulatory pathways related to morphological development. Finally, to scrutinize that the identified candidate genes do not just correspond to genes under strong purifying selection, which are expected to show an accelerated fixation of linked neutral variants, we investigated their Direction of Selection (DoS) scores, but found no excess of purifying selection compared to the genomic background (Figure S17, also when including *G. adriatica* Figure S18).

### Conclusion: The importance of considering genomic convergence at different molecular levels

Convergently evolved morphological traits often correspond to highly complex biological structures with a complex genetic basis expected to be highly polygenic (Cakan et al. 2012), involving gene interactions (epistasis) (Conith et al. 2018) and pleiotropic effects (Morris et al. 2019; Rennison and Peichel 2022). This is also expected for convergent traits in *Gouania*, such as the shape of the neurocranium or other complex morphological features, such as the number of vertebrae. Despite these difficulties, our analysis yielded evidence supporting convergent evolution at the level of single nucleotides, with identified genes strongly enriched for relevant developmental biological pathways involved in slender and stout morphotypes of *Gouania*. Furthermore, we found evidence of the involvement and regulation of transcription factors, such as *hox* genes, which are critical for bony structure development (Hubert and Wellik 2023).

In contrast, window-based analyses such as CSS – an established measure of parallel evolution – did not reveal any signatures of genomic convergence. This statistic, originally developed to study convergence in recently diverged stickleback populations, likely failed in our study, because of the relatively old age of the *Gouania* radiation (i.e., several million years versus thousands of years; (Marques et al. 2019; Wagner et al. 2019)) and concomitant old signals of genetic admixture.

This means that recombination had sufficient time in *Gouania* to unlink adaptive alleles inherited from a common ancestor from their genetic background.

Although we cannot conclusively address what proportion of convergently evolved single nucleotide variants are identical by descent (as opposed to corresponding to recurrent mutations), at least the signals of convergent evolution around *adam12* are not expected to have arisen independently. They rather point towards a single mutational origin of this putatively adaptive allele which further underlines the importance of old genetic variation in ecological diversification (Marques et al. 2019). Overall, our study highlights the importance of considering layered genomic convergence as it could be diagnostic of different underlying processes and timescales of convergent evolution.

## MATERIAL AND METHODS

### Diversity screening (DNA-barcoding analyses) and species delimitation methods

To assess the geographic variability and genetic diversity in the *Gouania* genus, we analysed a total set of 668 DNA barcodes (partial sequences of the mitochondrial cytochrome-c-oxidase subunit 1 (COI) gene) from 23 populations. A total list of all barcodes downloaded for this study (*Gouania* spp.: N=122 from (Wagner et al. 2017; Wagner et al. 2019); *Lepadogaster* ssp.: N=11 from (Wagner et al. 2017) and newly created barcodes (N=544, Genbank accession numbers OL839338-OL944591) can be found in Table S7). We generated the DNA barcodes following the procedures and protocols outlined in (Wagner et al. 2019) using the primers CO1_Fish_F1: 5’ TCAACCAACCACAAAGACATTGGCAC 3’ and CO1_Fish_R1: 5’ TAGACTTCTGGGTGGCCAAAGAATCA 3’ (Ward et al. 2005).

All sequences were aligned using MUSCLE (Edgar 2004: 20) in MEGA v7.0 (Kumar et al. 2016) and we tested for the best fitting substitution model according to BIC (K2+G). Then we conducted a phylogenetic analysis based on three datasets including all specimens (I), collapsed identical haplotypes (II) and species-specific datasets (III). For some of the analyses that require outgroups we used sequences of the genus *Lepadogaster* downloaded from GenBank. We used the collapsed dataset mainly for reducing the computational load for the Bayesian barcoding clustering analyses (see below). Then we calculated a maximum likelihood tree (ML) in IQtree v2.1.4-beta (Minh et al. 2020) using the automatic evolutionary model search function (Kalyaanamoorthy et al. 2017) and applying a resampling of 1000 ultrafast bootstraps (Hoang et al. 2018). Additionally, we calculated a maximum clade credibility tree (MCC-tree) for the reduced dataset in BEAST v2.6.6 (*50*), using a Birth-Death prior, applying the best model inferred in MEGA v7.0 (Kumar et al. 2016: 201) and a uniformly distributed substitution rate (0.0323 [0.022, 0.0424]; obtained from (Conway et al. 2017) for COI – see (Wagner et al. 2019)). The analysis was run for 200 million generations, with log- and tree-files stored every 20,000 generations. Finally, we assessed the convergence of the MCMC runs in Tracer v1.7.1 (Rambaut et al. 2014) and visualized the trees in FigTree v1.4.4 (available at http://tree.bio.ed.ac.uk/software/figtree/).

To assess species richness within the genus we employed a variety of single locus species-delimitation methods. In the first step we applied the distance-based DNA clustering algorithms Assemble Species by Automatic Partitioning (ASAP; (Puillandre et al. 2021)) and Automatic Barcode Gap Discovery (ABGD; (Puillandre et al. 2012)) by applying the Kimura-2-parameter model. We selected two thresholds 0.011 and 0.063 for ABGD because they represent the highest intraspecific divergence values in two plateau phases in the estimated number of groups (Figure S19). On the reduced haplotype dataset (II), we calculated the Bayesian Poisson Tree Processes (bPTP) model (Zhang et al. 2013), using specimens of *Lepadogaster* as an outgroup. We discarded a burn-in fraction of 0.25, ran the analysis for 5,000,000 MCMC iterations using a sampling frequency of 100 and visualised the values obtained from the heuristic and maximum likelihood searches. Furthermore, we applied the Mixed Yule Coalescent (GMYC) model (Fujisawa and Barraclough 2013) on the collapsed MCC tree using the “gmyc” function from the SPecies LImits by Threshold Statistics (splits) library in R.

### Classical and 3D geometric morphometrics

Based on 26 linear measurements obtained from (Wagner et al. 2021), we conducted a Principal Coordinates Analysis (PCoA) using the ape package (Paradis and Schliep 2019) based on a dissimilarity matrix using bray-curtis distance indices from the vegan package (Dixon 2003) in R v4.2.1 (R Core Team 2021). The individual measurements were standardized by standard length (SL). Additionally, we used 3D geometric morphometrics to compare cranial and pelvic girdle morphology across *Gouania*. Micro-CT scans representing the five *Gouania* species (n= 5-6 individuals per species) were obtained from a previous study (Wagner et al. 2021) and MorphoSource.org. We generated skeletal models of the neurocrania and basipterygia and placed landmarks on them using the SlicerMorph toolkit in 3D Slicer (Kikinis et al. 2014; Rolfe et al. 2021). 33 fixed landmarks were placed on each neurocranium, and 18 fixed landmarks and 6 semi-landmarks on each basipterygium. We performed a Generalized Procrustes Superimposition on each of the landmark data sets in R with the *geomorph* package (Adams and Otárola-Castillo 2013). To visualize the major axes of shape change, we conducted principal component analyses and plotted shape variation in morphospace. We warped representative neurocranial and pelvic girdle meshes to estimate shapes associated with extreme ends of the morphospace using the *Morpho* R package (Schlager 2017).

### Whole genome re-sequencing, draft-genome assemblies, and variant calling/filtering

We re-sequenced whole genomes of a total of 59 fish for this study based on Illumina short read 150bp paired end data. The obtained sequencing coverage ranged from 5 – 30X and we aimed for a balanced sample for each of the five *Gouania* species and morphotypes, “slender” and “stout” (i.e., 10-13 individuals per species). Additionally, we re-sequenced one specimen representing an outgroup of the genus, *Lepadogaster*. All raw data is stored at European Nucleotide Archive (ENA) under the project accession number PRJEB4981 (Table S8).

In addition to the existing *Gouania* reference genome (derived from a *G. adriatica* individual) we assembled draft genomes from our short-read data for the other four *Gouania* species and the outgroup species *Lepadogaster lepadogaster de novo*. For the genome assemblies we initially filtered raw reads with trimmomatic v0.39 (Bolger et al. 2014). Read quality for raw and filtered reads was assessed with FastQC v0.11.9 (Anders 2010) and summarised using MultiQC v1.9 (Ewels et al. 2016). Next, we used DISCOVAR de novo (Weisenfeld et al. 2014), SOAPdenovo v2.40 (Luo et al. 2012) and ABySS v2.0.2 (Simpson et al. 2009) to assemble filtered reads into contigs. Since ABySS requires the kmer size as input we assessed the optimal size using kmergenie 1.70 (Chikhi and Medvedev 2014) along a range of kmer sizes (21-121) with a step-size of 6.0. For each “best” obtained kmer size we added and subtracted a value of 4 and assembled contigs for each specimen using all three kmer sizes (referred to as “optimal k” = best k, “lower k” = best k - 4 and “upper k” = best k + 4) in ABySS. We assembled for each individual independently contigs in DISCOVAR de novo and SOAPdenovo. We assessed the quality and contiguity of the contigs in quast v5.0.2 (Gurevich et al. 2013). Contigs assembled from the above-mentioned three input kmer sizes were filtered according to highest N50 values (Table S1). Finally, the selected draft assemblies were scaffolded using a reference guided approach in RagTag v1.0.1 (Alonge et al. 2022) using the *Gouania* v. fGouWil2.1 (Rhie et al., 2021) reference genome as backbone. For assessing the “best” scaffolded assemblies we ran BUSCO v4.1.3 (Waterhouse et al. 2018) using the set actinopterygii_odb10. We selected draft-genomes with (first) the highest number of single copy BUSCO genes, (second) highest N50, and (third) lowest numbers of N and for the subsequent phylogenetic analyses (Table S2).

We then aligned raw reads from re-sequencing data of 59 fishes (10-13 individuals per species) against fGouWil2.1 (Rhie et al. 2021) using BWA v0.7.17 (Li and Durbin 2009) to create a call-set for further genomic exploration. After inspection of the mapping quality using samtools flagstat, we obtained variant sites using bcftools v1.14 (Danecek et al. 2021). We filtered/masked sites which had a mapping quality of zero for more than 10% of reads mapping to that site or which showed an overall mapping quality of less than 50. Additionally, we masked sites where the mapping quality between the two strands was significantly different (p < 0.001) and sites that showed extremely high (*>*97.5 percentile) or low (<2.5 percentile) depth. We filtered heterozygous sites which had a significantly biased depth of reference and alternative alleles by means of binomial test (PHRED score >20), which represents between 0.02% and 0.15% of heterozygous sites per individual. Furthermore, we masked sites showing InbreedingCoeff < 0.2 (i.e., excess heterozygosity) and sites with more than 20% of missing genotypes. For the subsequent investigation only biallelic SNPs or indels were considered leading to a total of 22,348,287 sites across the 23 chromosomes. Finally, we annotated the single variants and their effects using SnpEff (Cingolani, Platts, et al. 2012) and the gene annotation version fGouWil2.1.99.

### Phylogenomic analyses

For inferring the phylogenetic relationships within the genus *Gouania* we used the phylogenomic pipeline phylociraptor (commit #6a6c0eb) (Resl and Hahn 2023). Phylociraptor extracts BUSCO genes from (draft) genomes and calculates a concatenated maximum likelihood tree in IQtree v2.0.7 (Minh et al. 2020), a neighbour-joining tree and a species tree in Astral v5.7.1 (Zhang et al. 2018) based on the corresponding set of single copy BUSCO genes. In addition to our draft assemblies above, representing the species *G. willdenowi*, *G. pigra*, *G. hofrichteri*, *G. orientalis* and *L. lepadogaster*, we downloaded the latest reference genomes of *G. adriatica* (fGouWil2.2) and another outgroup of the family of Blenniidae, *Salarias fasciatus* (fSalaFa1.1), from NCBI (Rhie et al. 2021). Orthologous genes were identified with phylociraptor using BUSCO v3.0.2 and the gene set actinopterygii_odb9. We filtered out multicopy orthologs, orthologs with missing data in more than one individual (minsp=6) and less than 11 parsimony sites to retain only phylogenetically informative single copy orthologs. Alignment and trimming of the individual sequences for each locus was done using MAFFT v7.464 (Katoh and Standley 2013) and trimAl v1.4.1 (Capella-Gutiérrez et al. 2009). We performed phylogenomic inference on the remaining trimmed alignments of 3406 loci. For each locus we applied an automatic search for the best substitution model in IQtree (Kalyaanamoorthy et al. 2017) with 1,000 ultrafast bootstrap resampling replicates (Hoang et al. 2018)

From the concatenated data, we inferred a maximum-likelihood tree in IQtree with 1,000 ultrafast bootstrap replicates and a species tree in Astral using the default parameters in phylociraptor. Furthermore, for the tree based on concatenated data we computed site and gene wide concordance factors (Mo et al. 2023). We then tested three different constrained tree topologies: (i) a topology without convergence (i.e., two main clusters comprised of slender and stout morphotypes); (ii) *G. willdenowi* as sister to all other *Gouania* species (as a scenario explaining a refilling from the Mediterranean after the Messinian salinity crisis); and (iii) *G. hofrichteri* as the global outgroup (i.e., the neighbour joining topology). We compared them with an approximately unbiased (AU) test of phylogenetic tree selection (Shimodaira 2002) using 10,000 RELL replications (Kishino et al. 1990) in IQtree. We then calculated the gene-wise phylogenetic signal (dGLS) (Shen et al. 2017) to assess the contribution of single loci to the overall phylogeny.

Finally, to test for potential introgression events we constructed an explicit phylogenetic network from all trees inferred for all 3406 loci with the SNaQ (Species Networks applying Quartets) method (Solís-Lemus and Ané 2016). We used the Phylonetworks v.0.14 (Solís-Lemus et al. 2017), implemented in the Julia package using the code and procedures following (Lescroart et al. 2023).

### Window and site-wise investigation for searching candidate genomic regions

We conducted several analyses in defined window length along the genome to detect regions under convergent differentiation between groups. For the window-based analyses we excluded indels, but not for the site-based analyses (see below).

In a first step, we used the “popgenWindows.py” script from https://github.com/simonhmartin/genomics_general to estimate the genetic diversity (*pi*) of each of the five *Gouania* species and to calculate average net between-group divergence (*D_a_*):

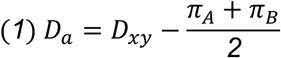

where *D_xy_* is the average genetic distance between two species and π_A_ and π_B_ represent the intraspecific diversity within each species A and B. We calculated *D_a_* in sliding windows of 20,000 bp and a step size of 10,000 bp for each of the 23 chromosomes for all species combinations. Furthermore, we calculated the cluster separation score (CSS;(Jones et al. 2012; Miller et al. 2019)) between the slender and stout *Gouania* morphotypes, mainly following the procedures described in (De-Kayne et al. 2022) using a window size of 20,000. For *D_a_* in windows, we considered as outliers the upper five percent of all data points and for CSS a permutation test was applied as in (De-Kayne et al. 2022). We then extracted all gene information within the window boundaries of outlier coordinates using SnpSift (Cingolani, Patel, et al. 2012).

### Site-wise investigation for searching candidate genomic regions

We assessed the extent of convergent evolution on single sites. We, firstly, estimated the allele frequency at each site for each of the five *Gouania* species using bcftools. Then, we developed an allele frequency-based statistic to assess the extent of convergent evolution for two sets of groups (in our case *slender* and *stout*) belonging to four species using the formula below:

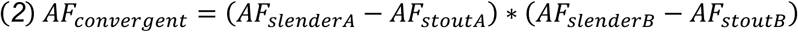

AF corresponds to the allele frequency and A and B to the two convergent slender and stout species pairs. In our concrete case we tested the species *G. hofrichteri* (slenderA) and *G. orientalis* (stoutA) as well as *G. pigra* (slenderB) and *G. willdenowi* (stoutB). We chose this species combination because *G. orientalis* is the sister species of *G. adriatica* and therefore would not add independent genetic information in terms of convergent evolution in the genus. We assume that all sites of convergence scores that are smaller than -0.25 or larger than 0.25 to be outliers since they correspond to an average allele frequency change of 0.5 between the morphotypes.

Since we found an interesting region with many positive outliers in and around the gene *adam12* on chromosome 15 (position 10,900,000–11,111,000), we further investigated this region in greater detail. We also extracted an IUPAC coded multiple sequence alignment from vcf files of exon 3 of *adam12* and visualized the translated sequence based on the reverse strand (the direction of expression of *adam12*) as sequence logos using WebLogo3 (Crooks et al. 2004).

Then, to test for convergent fixed mutations identical by state, we checked for alleles that were fixed in either stout or slender morphotypes but not between any stout or slender. Specifically, we checked for SNPs or indels that are fixed for one allele in either of the *Gouania* species but fixed for the other allele in all other species (fixed differences). We then extracted a list of genes associated with these variants from the SNPeff annotation and removed duplicated gene names. Next, we checked for candidate genes using the formulas below (note that the all species comparison is only available in the Supplementary Results/Discussion).

*A* two-species comparison (excluding *G. adriatica*):

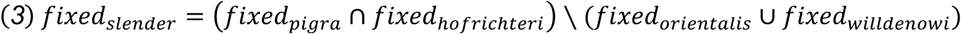

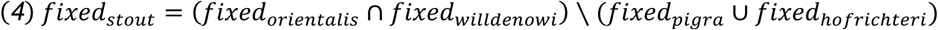

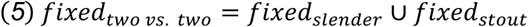

and *B* all-species comparison:

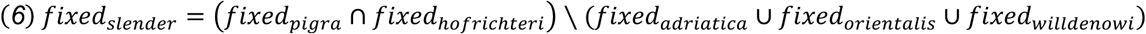

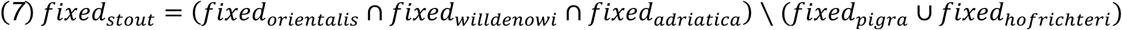

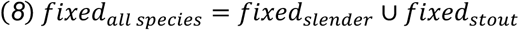

Since fixed differences might reflect regions under strong purifying and not divergent selection, due to a lower effective population size and increased drift in these regions, we did a McDonald-Kreitman test (McDonald and Kreitman 1991) based on the SNPeff annotation. Because mean NI values can be biased, we calculated genome-wide NI_TG_ and to test whether NI_TG_ values significantly differed from 1 (the expectation under neutrality), we performed a parametric bootstrap with 1,000 iterations. We then calculated the direction of selection (DoS) for single genes following (Stoletzki and Eyre-Walker 2011) and applied a Fisher’s exact test to determine significant positive selection (i.e., NI < 1) from all available genes and species pairs.

To assess potential functional convergence, we performed gene ontology (GO) enrichment analyses using g:Profiler (Reimand et al. 2007) on all gene sets identified in the convergence score and fixed allele analyses. All the GO analyses were run in the same way using the candidate gene lists as input, zebrafish as organism, a custom background gene list (i.e., all genes obtained from the snpeff annotation) and applying a significance threshold of 0.05 based on Benjamini-Hochberg FDR (Benjamini and Hochberg 1995).

For the candidate genes obtained from the fixed AF differences, we furthermore randomly reshuffled the formulas (3 and 6) above and conducted an independent GO analyses for each obtained gene set. Then we checked for overlaps and unique GO terms of the random sets and the obtained biologically meaningful (slender vs. stout) comparisons. Finally, we also checked for genes enriched for certain transcription factor binding sites (TFBS) from the input gene sets in g:Profiler following the same procedure described above for the GO analysis.

## Supporting information

Supplementary Results/Discussion and Figures

## ACKNOWLEDGEMENT/FUNDING

The authors thank NOBIS-Austria for the financial support (NOBIS-Scholarship to M.W.). M.W. has also been supported with a DOC-Scholarship by the Austrian Academy of Science during his PhD and a YUFE4Postdoc-Scholarship, co-funded by the EU under Marie Sklodowska-Curie Grant Agreement No. 101081327. Finally, we also want to thank Sandra Bračun, Gerhard Skofitsch, Marcelo Kovačić, Samuel P. Iglésias and Stamatis Zogaris for their support throughout the last years, during sampling and for the creation of the MicroCT images.

## DATA AVAILABILITY STATEMENT

The newly generated DNA barcode sequences, raw-sequencing files and callsets underlying this article are available in GenBank Nucleotide Database (https://www.ncbi.nlm.nih.gov/; Accession Numbers OL839338-OL944591), European Nucleotide Archive (ENA; https://www.ebi.ac.uk/ena/browser/home; Project accession number PRJEB49819) as well as European Variation Archive (EVA; https://www.ebi.ac.uk/eva/; Project accession number PRJEB60875). Individual accession numbers are listed in the supplementary material. The bioinformatic pipelines, written in snakemake (Mölder et al. 2021), and the jupyter notebook (used for local analyses) can be accessed under https://github.com/maxwagn/Gouania-Phylogenomics.

## AUTHOR CONTRIBUTIONS

Conceptualization: MW, SK, HS, RD; Methodology: MW, HS, RD, MS, IB, SM, NK, PR; Investigation: MW, PR, NK, JMH, IB, SM, HS; Visualization: MW, PR, NK, JMH; Supervision: RD, SK, HS; Writing—original draft: MW, HS, SK, PR, JMH; Writing—review & editing: MW, HS, SK, PR, JMH, IB

## Notes

### Competing Interest Statement

The authors have declared no competing interest.

### Summary of Updates

In the original post, the equations were not displayed correctly due to a conversion error.

