## Supplementary Results/Discussion and Figures for "The genomics of convergent adaptation to intertidal gravel beaches in Mediterranean clingfishes"

Maximilian Wagner *et al.*

### SUPPLEMENTARY RESULTS/DISCUSSION

#### DNA Barcoding reveals that a five species model is most plausible

Cryptobenthic fishes are prone to showing high levels of cryptic diversity (i.e., species that are considered morphologically identical; reviewed by (Brandl et al. 2018)). Being no exception, the blunt-snouted clingfishes of the Mediterranean Sea were long considered to be a single species before a recent taxonomic revision established five morphologically distinct species (Wagner et al. 2019; Wagner et al. 2021). Therefore, we sought to find potential cryptic or overlooked biodiversity within the blunt-snouted clingfishes, which could lead to misinterpretations in the scope of phenotypic convergence.

We analysed 668 mitochondrial cytochrome-c-oxidase I (COI) DNA barcodes from 23 populations of all described species across previously recorded sites (Figure S3). Our analyses show differences between different barcoding algorithms: while distance-based species delimitation methods (ASAP; ABGD) seem compatible with the current taxonomic situation (i.e., five species), tree-based methods (PTP; GMYC) overestimated the current five species taxonomic consensus (Figure S3a). Generally, tree-based species delimitation methods are prone to over-splitting, and can be influenced by haplotype-richness, the sampling design, effective population size and the presence of singletons (summarized by (Guo and Kong 2022)). Additionally, evidence of clear geographical (Figure S3b; except for *G. orientalis* populations from the Ionian Sea) or morphological splitting of populations was lacking (compare with (Wagner et al. 2021)).

We therefore conclude for this study that potential cryptic diversity will not alter our interpretation of phenotypic convergent evolution within the genus. However, future studies incorporating samples from currently unexplored regions, such as the Middle East or the North African coastline, could uncover even greater diversity and may necessitate a reassessment of convergence within the genus  (Wagner et al. 2021).

**Alternative phylogenetic hypotheses testing confirms that Gouania morphotypes evolved convergently**

Especially the low node support values towards the root could impact our interpretation of convergent phenotypic evolution in *Gouania*. Indeed, we found evidence of 38% introgression from the ancestor of *G. hofrichteri* into the ancestor of *G. orientalis* and *G. adriatica*, which could explain this measured inconsistency (Figure 2d). Thus, before investigating the genomic basis of convergently evolved phenotypes, we wanted to exclude any other (non-convergent) alternative phylogenetic scenario.

We, therefore, tested the likelihood of three alternative phylogenetic scenarios (Figure S6): i) a topology without convergence (i.e., two main clusters comprised of slender and stout morphotypes); ii) *G. willdenowi* as sister to all other *Gouania* species (as a scenario explaining a refilling from the Mediterranean after the Messinian salinity crisis); iii) *G. hofrichteri* as the global outgroup (i.e., the neighbour joining topology; Figure S5). Overall, we found lower support for these three alternative topologies compared to the best tree (Figure 2a). Consistent with this, an approximately unbiased (AU) test of phylogenetic tree selection was non-significant for all three alternative topologies (Shimodaira 2002) (Table S4).

We then quantified the fraction of genes which contributed to alternative topologies, and calculated the gene-wise phylogenetic signal (dGLS; (Shen et al. 2017).  We found that, from all three alternative topologies, the topology with slender *G. hofrichteri* as an outgroup to all other *Gouania* (iii) was supported by 42% (1437 of 3,406) of loci (Figure S6). On the contrary, the least supported one, was the one which described a non-convergent scenario (i), as it only had negative dGLS values in 26% (915 of 3,406) of loci (Figure S6). Consequently, we conclude that convergent phenotypic evolution is the most likely phylogenetic scenario in our *Gouania* dataset.

**Including Gouania adriatica in the pairwise allele-frequency comparison does elevate the number of overall loci but results remain similar**

For reasons of clarity and to stay with a two versus two study design we removed *Gouania adriatica* from the analysis including independently fixed alleles (Figure 4). However, we conducted the same analysis again for the whole dataset. Overall, by including *G. adriatica* in the pairwise allele-frequency comparison we increased the number of overall loci to 2,011 (as compared to the analysis without *G. adriatica*).

For *G. pigra* we found 13,294 variants associated with 7,136 genes, for *G. hofrichteri* 70,438 variants associated with 13,189 genes, for *G. adriatica* 1,113 variants associated with 946 genes, for *G. orientalis* 1,062 variants associated with 918 genes, and for *G. willdenowi* 13,724 variants associated with 7,208 genes. Whereas 2,010 associated genes are shared between the two slender morphotypes, but not present in any of the stout morphs, only one associated gene was found in all three stout morphs but was not present in any of the slender morphs (Figure S16a). This discrepancy can be explained by the nature of the experimental design (three stout vs. two slender comparison) and the phylogenetic relationships of these species. For the total of 2,011 genes, we detected 35 significantly enriched biological process categories (Figure S16b). Like for the reduced dataset (Figure S17), we could not detect an excess of purifying selection for these 2,011 genes (Figure S18). Next, to exclude categories not specifically enriched for parallel evolutionary processes, we repeated the same test for all possible combinations of species by randomly permuting stout and slender morphotype labels, thereby testing for enrichment of genes with fixed differences for non-parallel scenarios. We found that 14 out of the original 35 GO terms were uniquely enriched in the parallel comparisons (bold terms in Figure S16b). These GO terms include skeletal system morphogenesis, embryo development ending in birth or egg hatching or chordate embryonic development and are therefore, overall similar compared to the dataset excluding *G. adriatica* (Figure 4d).

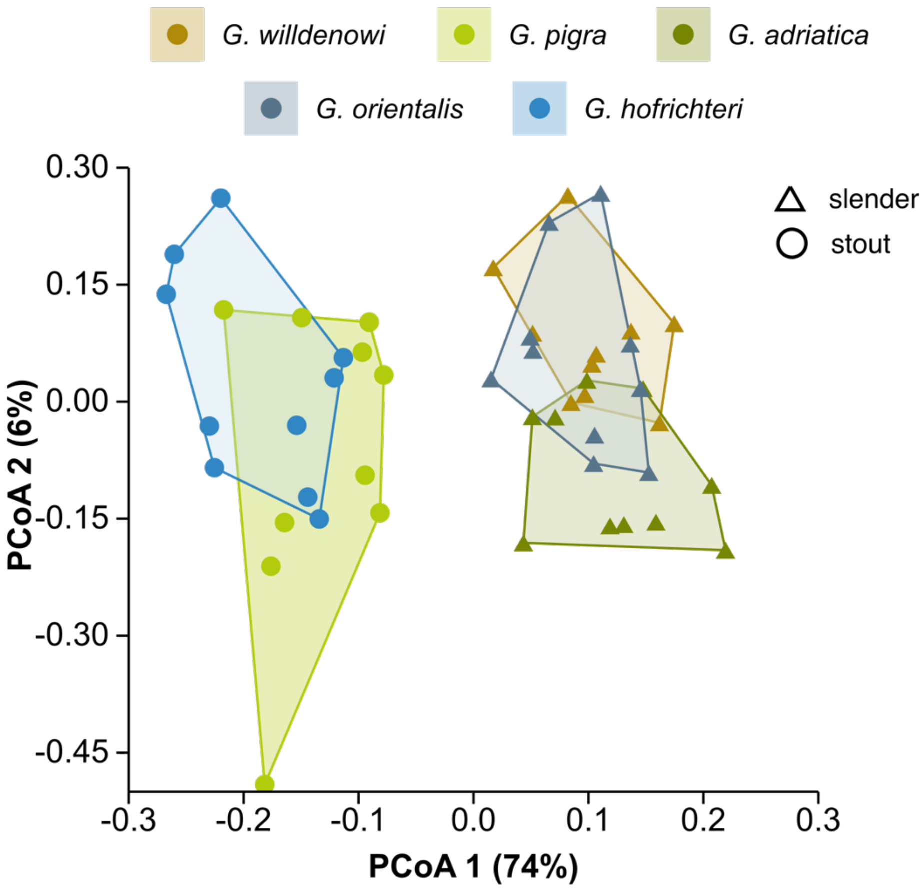


**Figure S1.**

Principal Coordinates Analysis (PCoA) of 26 linear body measurements (using the data in (Wagner et al. 2021)) based on dissimilarity matrix using Bray-Curtis distance indices.

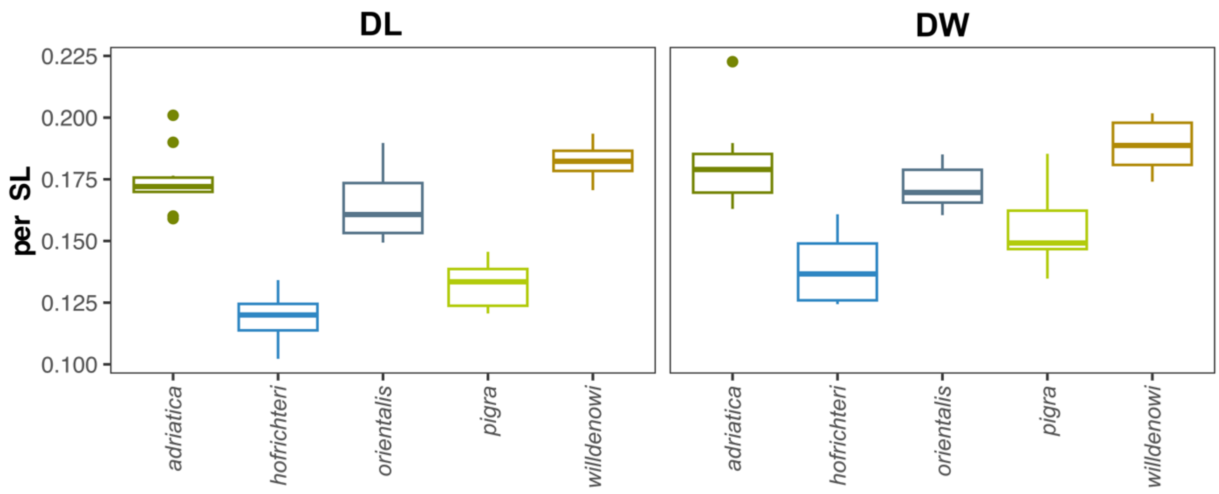


**Figure S2.**

Measurements of the ventral sucking disc length (DL) and disc width (DW) across different *Gouania* species.

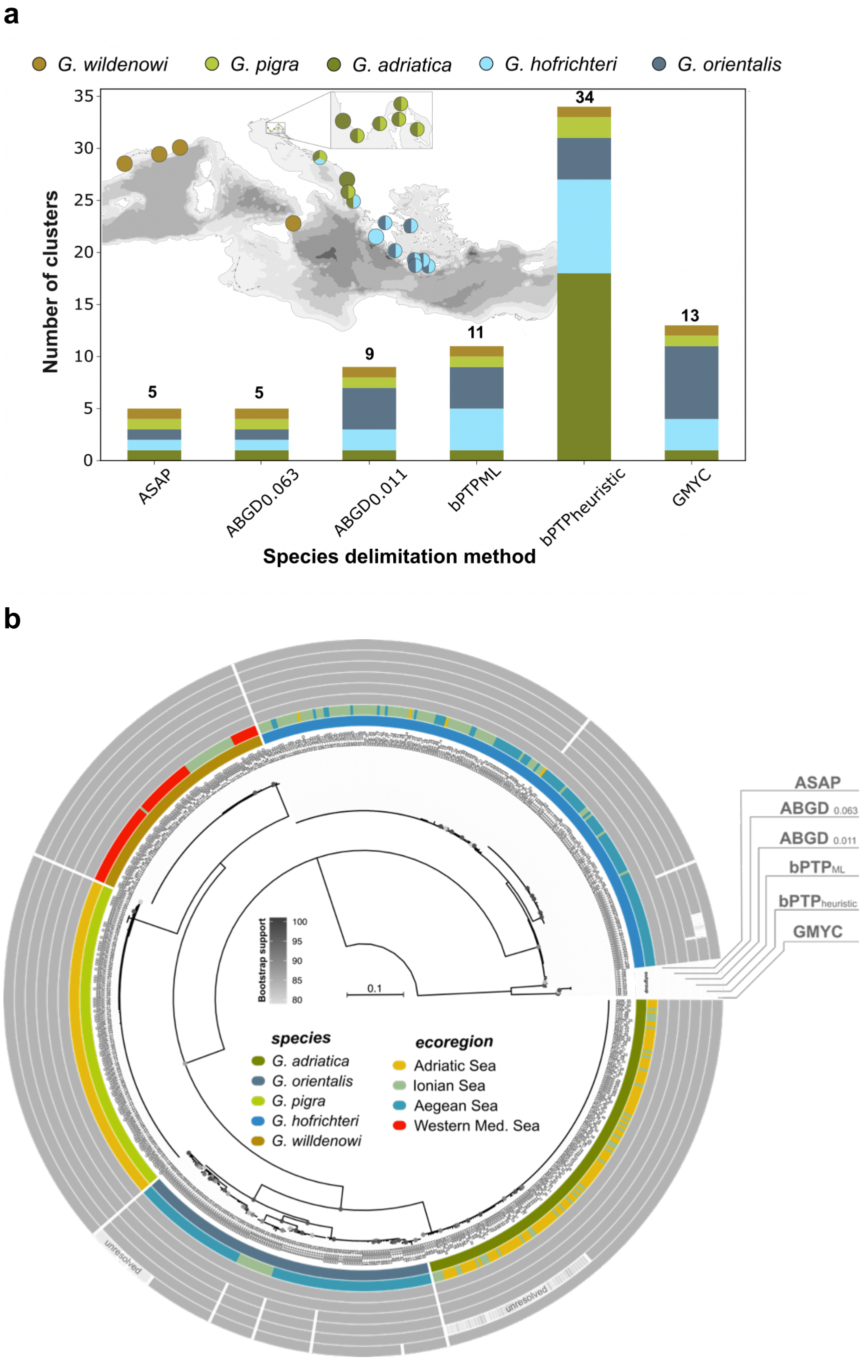


**Figure S3.**
DNA Barcoding and species delimitation methods summary (a) Maximum likelihood-based COI barcoding tree (IQtree) annotated according to different clusters obtained by species delimitation methods. (b) Overview of the number of clusters obtained from different distance-based (ASAP, ABGD) and tree-based (bPTP, GMYC) molecular species delimitation methods. Samples were obtained from 23 sites across the distribution ranges from all known *Gouania* species.

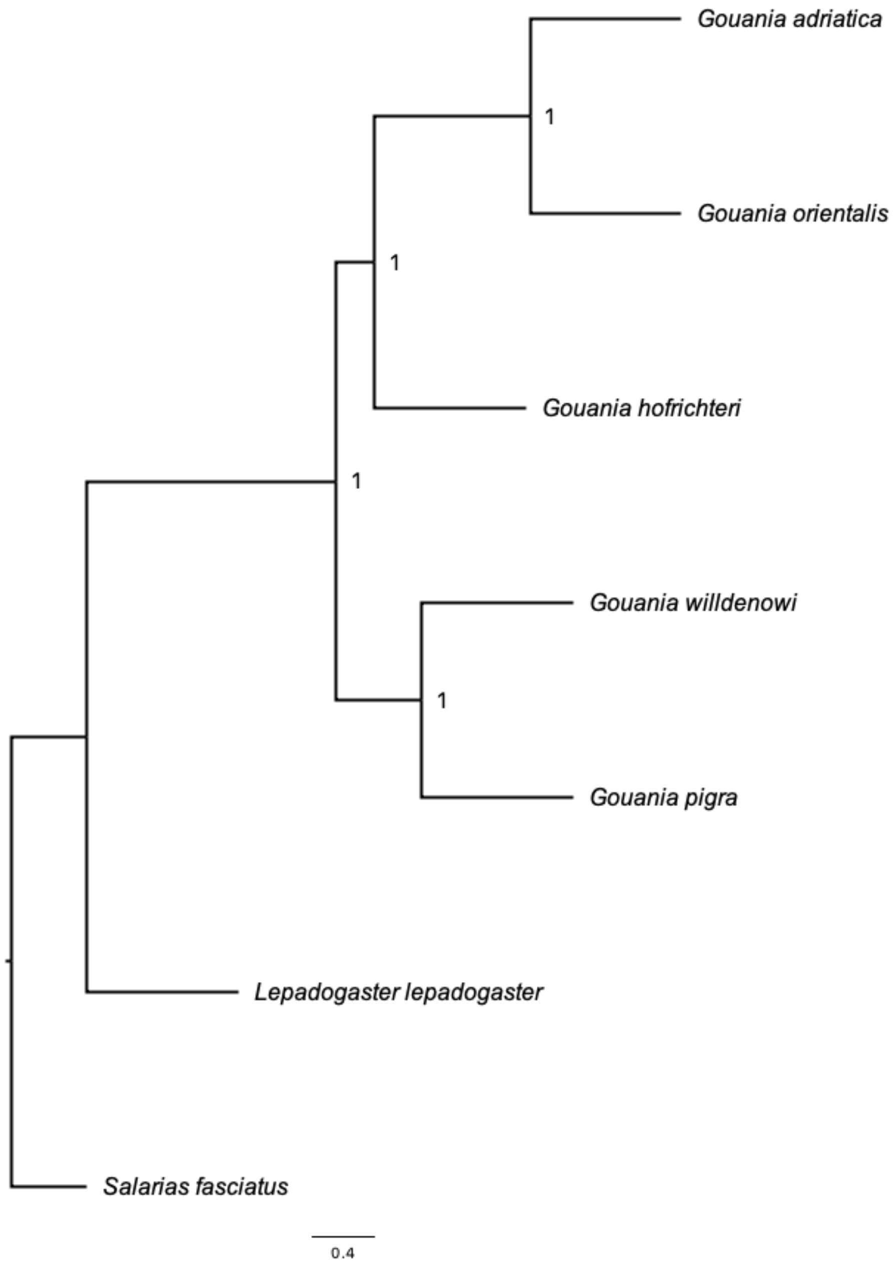


**Figure S4.**
Astral species tree based on 3406 single copy orthologous BUSCO genes. For each node the quartet score is given.

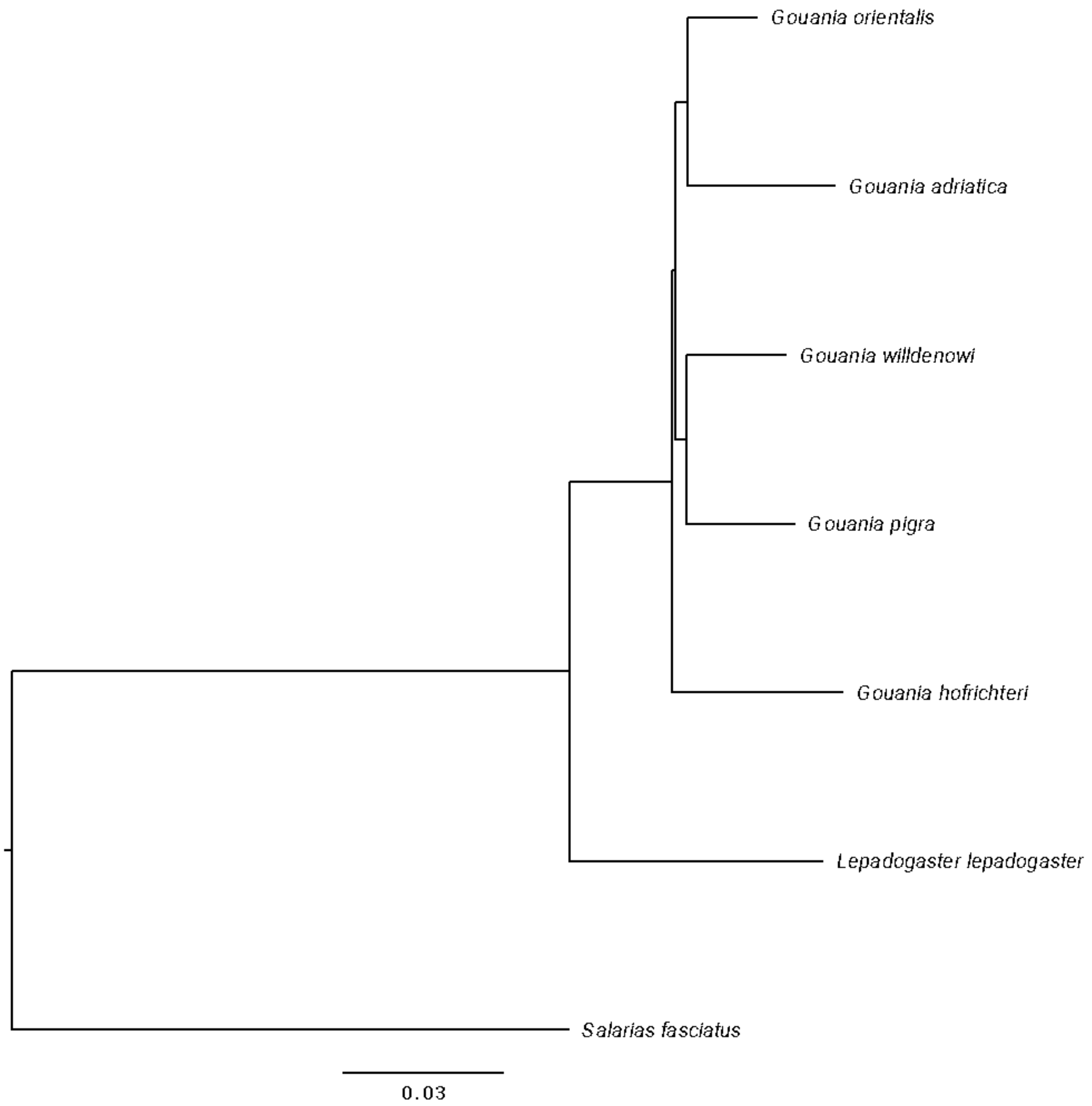


**Figure S5.**
Neighbor-Joining Tree based on a distance matrix from a concatenated alignment of 3406 single copy orthologous BUSCO genes.

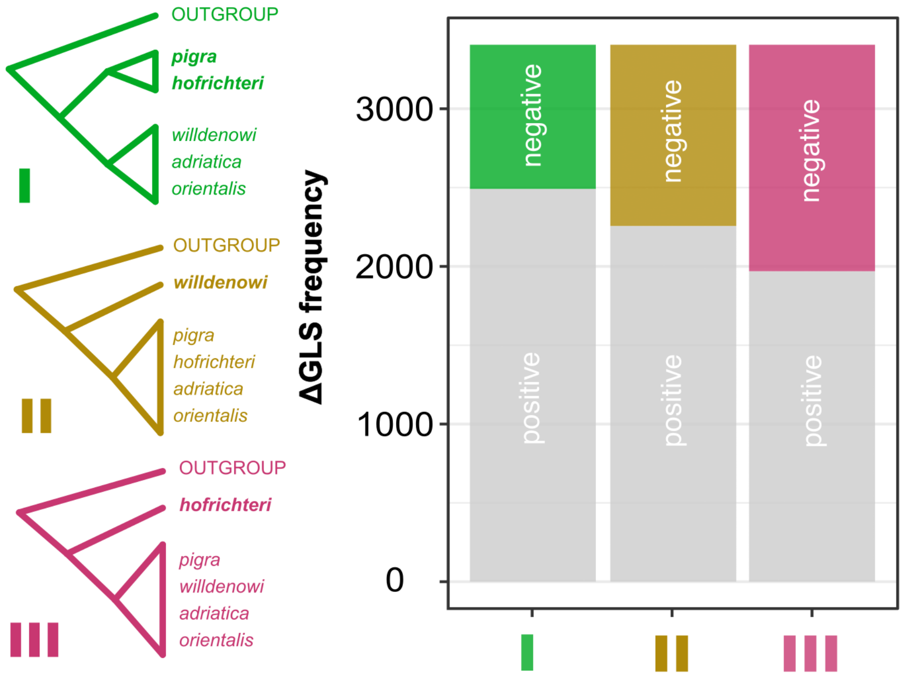


**Figure S6.**
Investigation of the gene-wise phylogenetic signal (dGLS) based on constrained trees obtained from 3406 single copy orthologous BUSCO genes. A negative dGLS value indicates support for the alternatively tested constrained topology (I to III), suggesting that a least likely scenario includes non-convergent evolution (I) followed by a western Mediterranean, *G. willdenowi*, ancestry (II) and the most likely of all three tested hypothesis is the Neighbour-Joining topology (G. hofrichteri as a global outgroup; III).

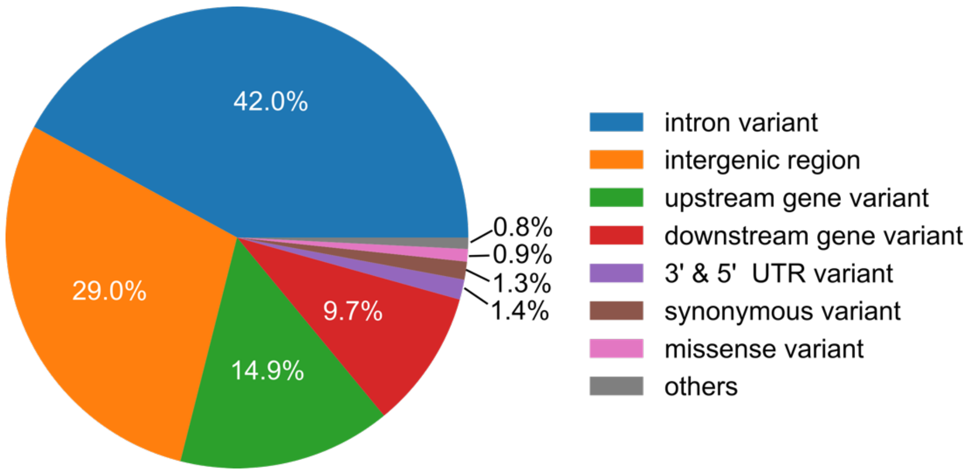


**Figure S7.**
The final variant call-set included 22,348,287 sites representing 18.,936,907 biallelic SNPs and 3,411,380 indels.

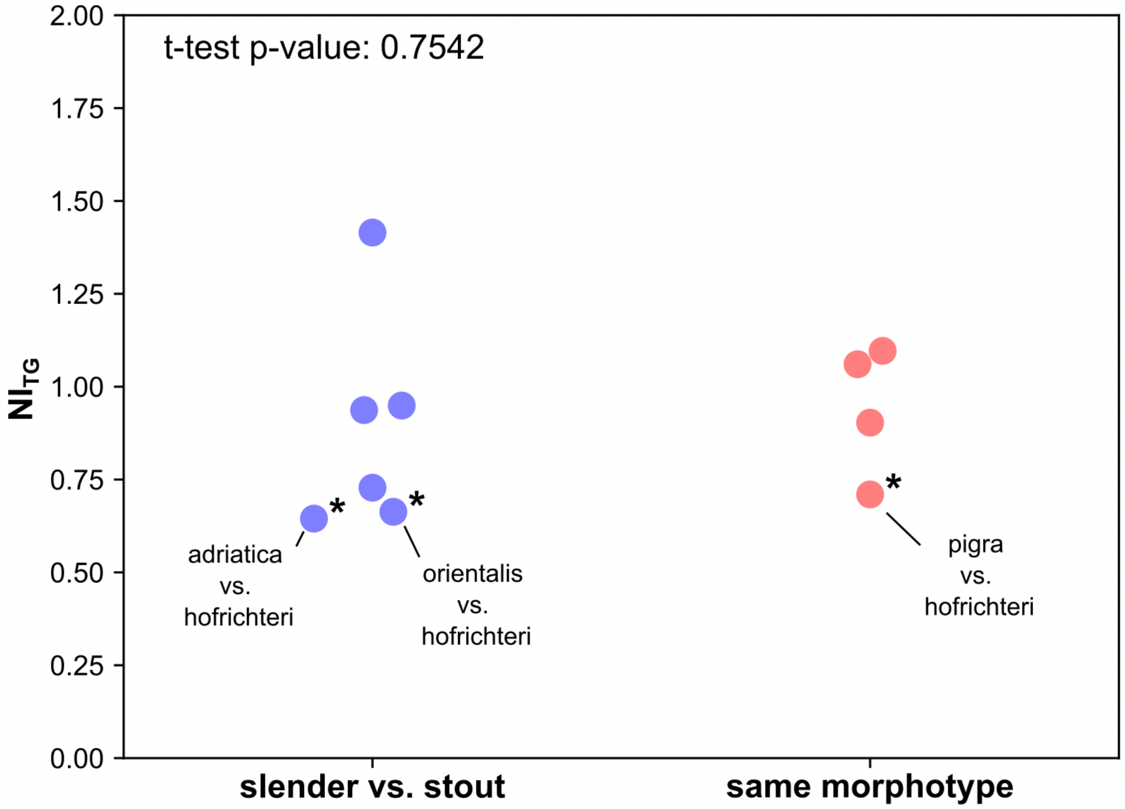


**Figure S8.**
Genome-wide pairwise signal of positive selection ($NI_{TG}$) shows no significant difference between slender and stout comparisons against same morphotype comparisons. Indicated with asterisks are $NI_{TG}$ values significantly smaller than 1, suggesting positive selection. P-values were obtained from 1000 parametric bootstrap samples.

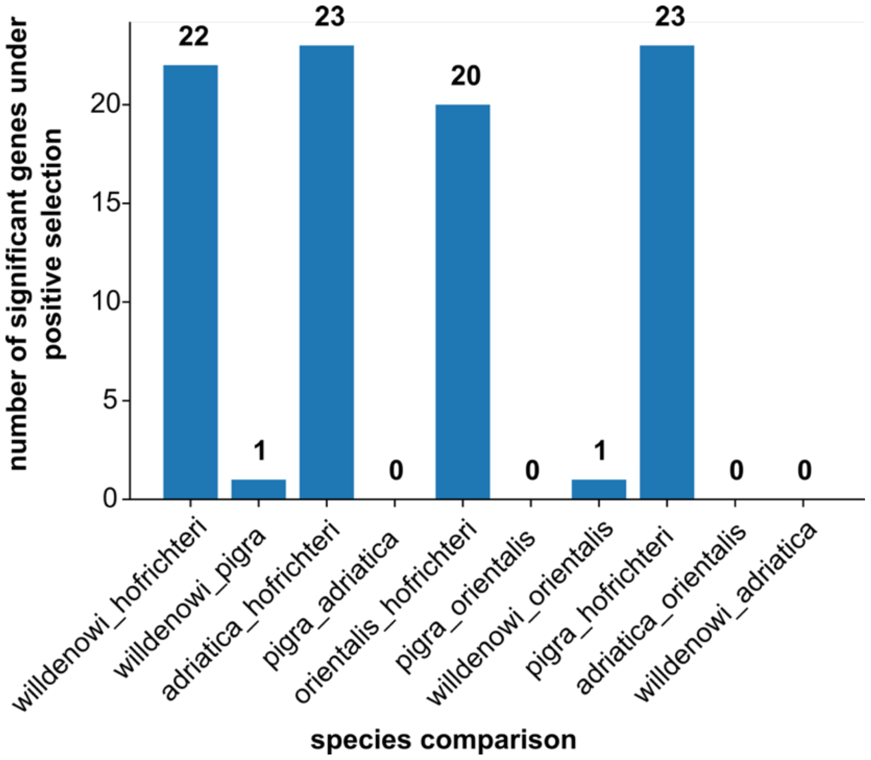


**Figure S9.**
Overall number of genes that are under significant positive selection (NI < 1; Fisher's exact test $p < 0.05$).

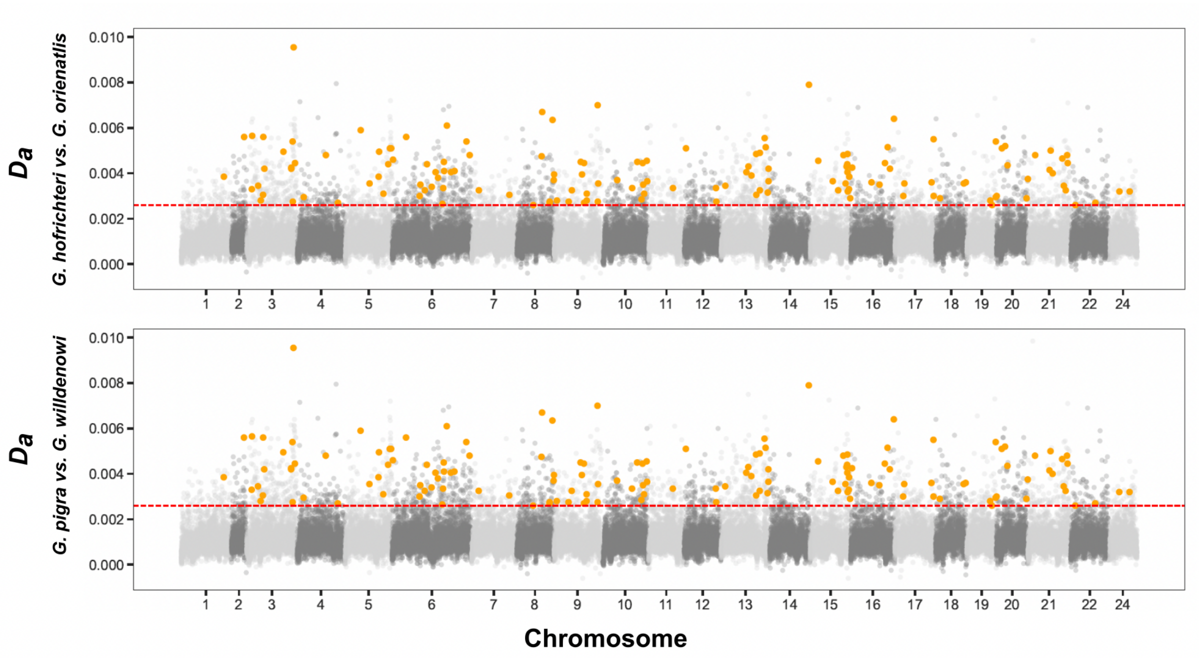


**Figure S10.**
$D_a$ (net between group mean divergence) values calculated in 20.000 bp windows along the genome. Highlighted in orange are dots that are shared between the two species comparisons. Red dashed lines indicate significant upper 5 % outliers.

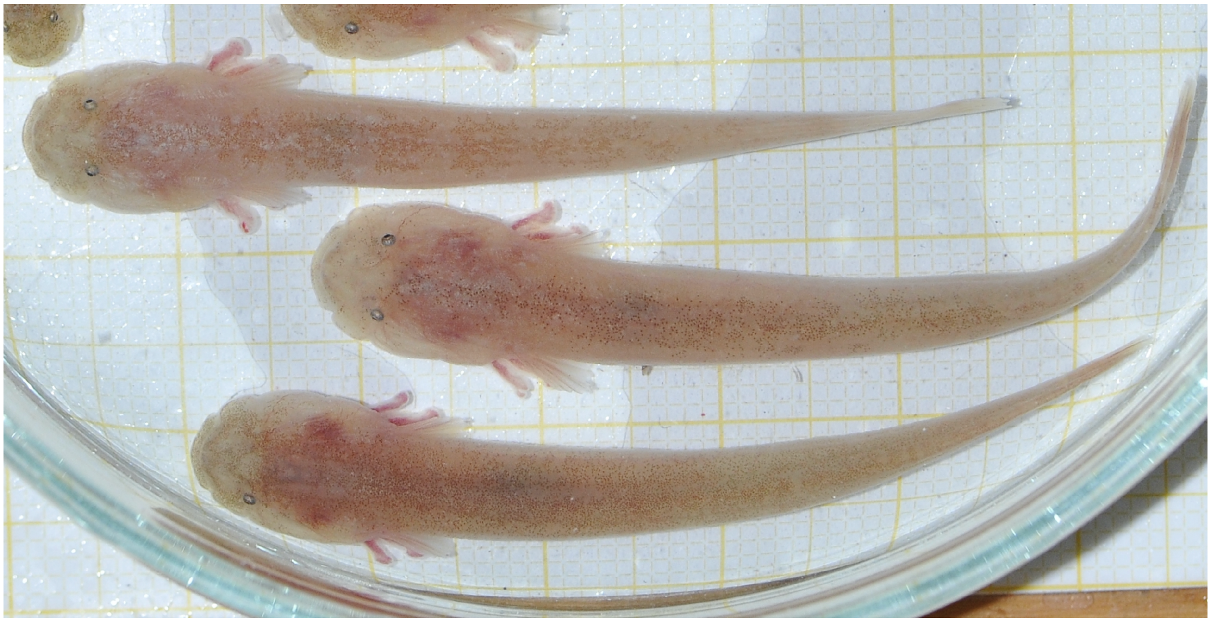

**Figure S11.**
Injured mature male missing the right eye (below) compared to two males with well-developed eyes.

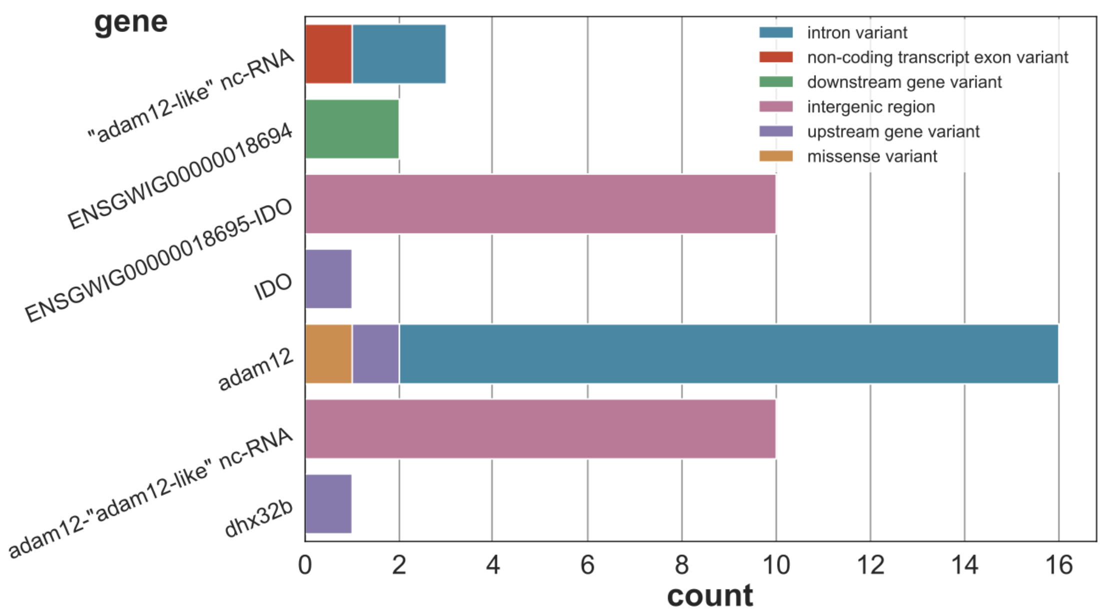


**Figure S12.**
Variant types within the "*adam12*-haplotype" on chromosome 15 are mainly present as non-coding changes.

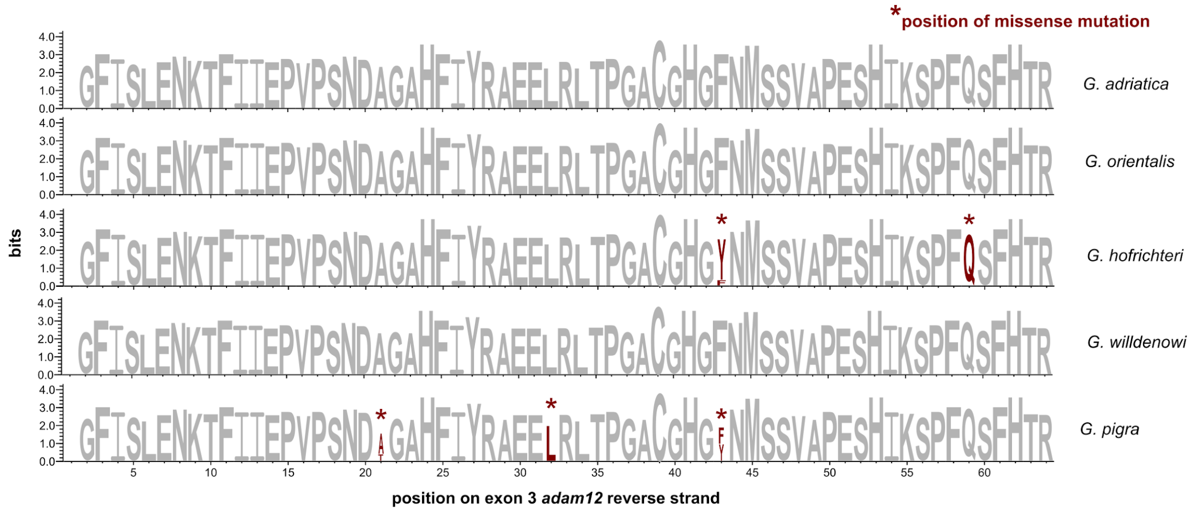


**Figure S13.**
Amino acid (AA) sequence of exon 3 of adam12 (reverse strand) showing relative contribution of each AA at certain mutations and missense mutations are highlighted by asterisks. The missense mutation on position 43 follows a convergent pattern according to the parallel score (Figure 3).

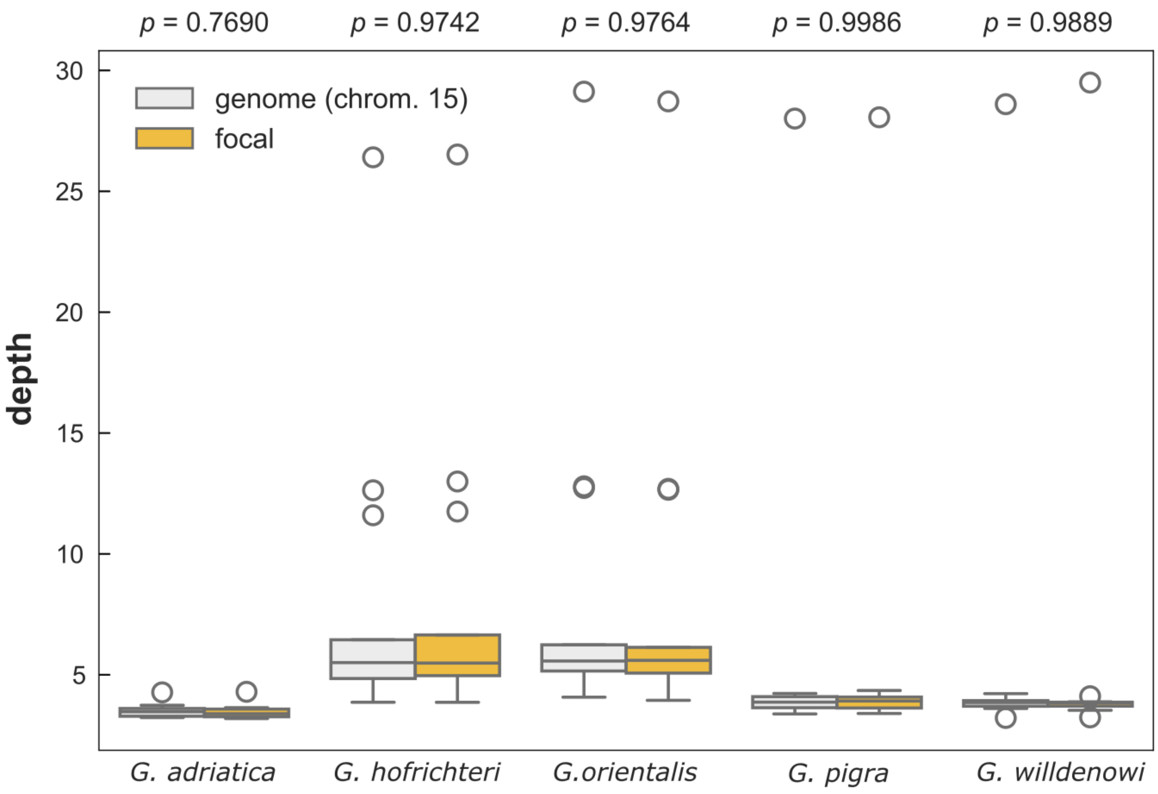


**Figure S14.**
There is no difference between the sequencing depth in the focal region ("*adam12*-haplotype" on chromosome 15) and the whole chromosome 15, which suggests that the variation in the region is not the result of a duplication event.

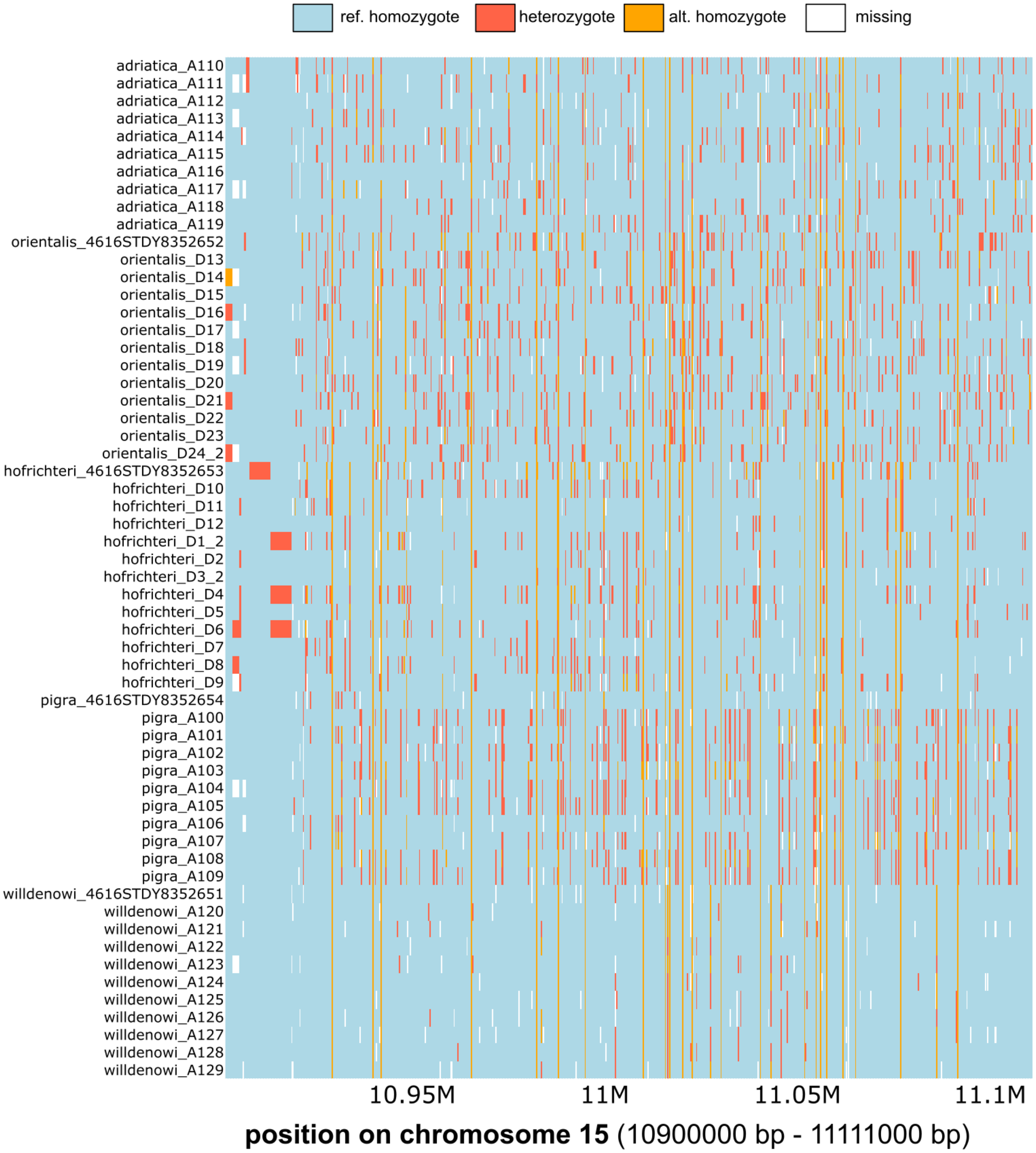


**Figure S15.**
Genotype plot showing the ”*adam12*-haplotype” on chromosome 15 shows that no continuous haplotype is present.

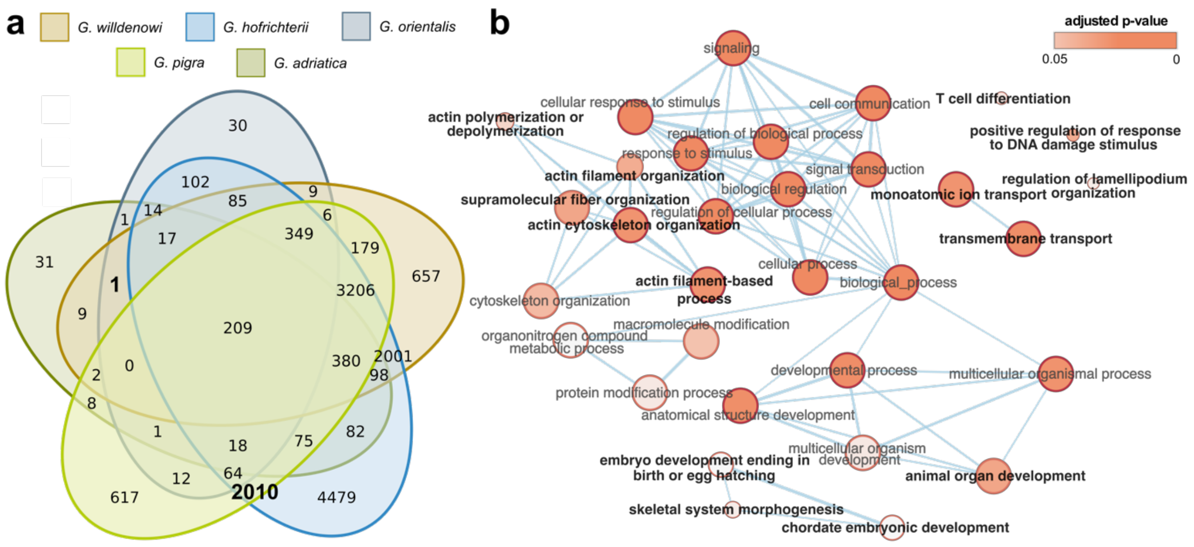


**Figure S16.**
Fixed differences associated with parallel patterns in the radiation of the blunt-snouted clingfishes including G. adriatica. (a) Venn diagram of numbers of genes containing fixed SNP differences between a species and the four other *Gouania* species. The categories corresponding to genes with differentially fixed SNPs only in the two stout species and only in the three slender species contain 2010 and 1 genes, respectively (in bold). (b) GO analyses showing biological processes enriched for the 2011 genes associated with parallel fixed variants (see a). Highlighted in bold are terms unique to parallel patterns in the genus after excluding terms which also occur in random (non-parallel) gene sets.

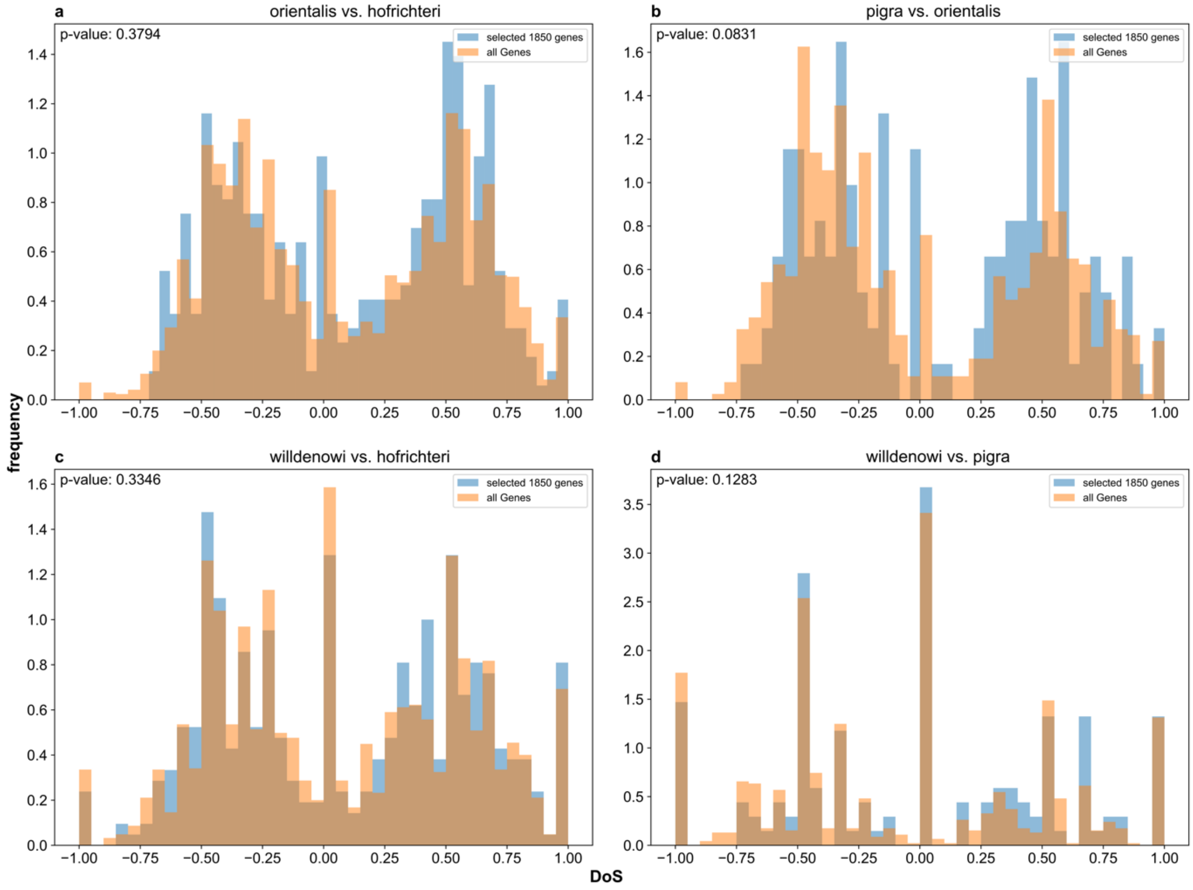


**Figure S17.**
No excess of purifying selection by means of Direction of Selection (DoS) values for the selected 1,850 selected genes for all slender versus stout comparisons (a-d) compared to the genomic background.

a

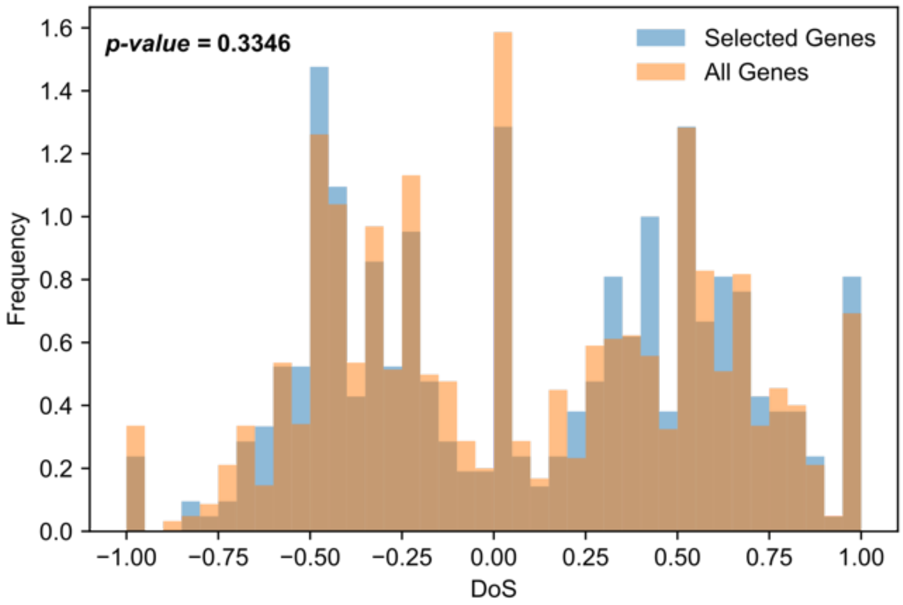


b

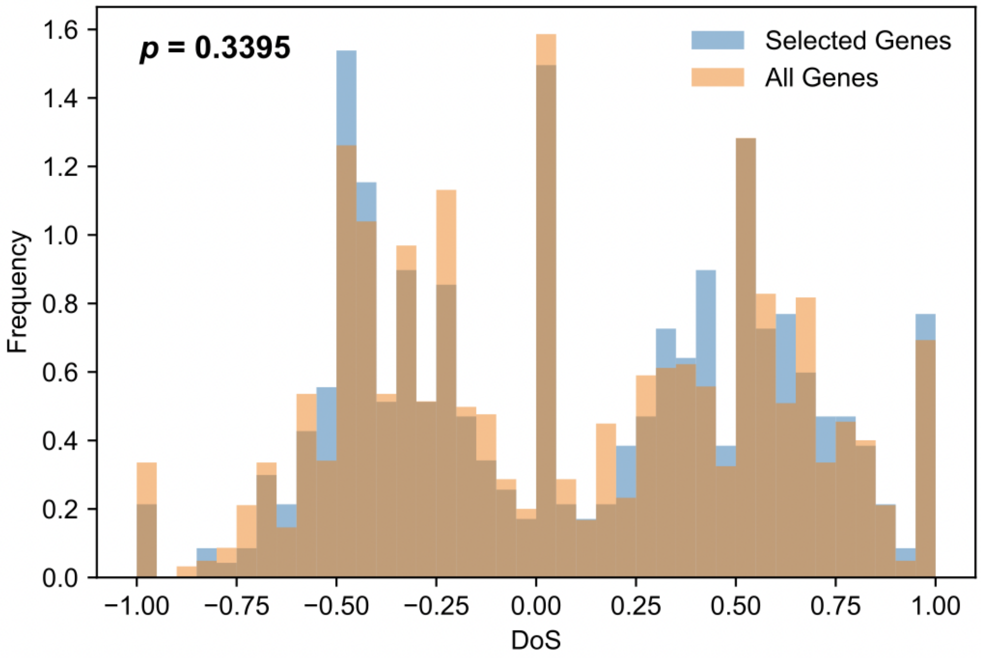


**Figure S18.**
Direction of Selection (DoS) scores based on the 1,850 (a; dataset without *G. adriatica*) and 2010 (b; full dataset – see Figure S8) selected genes unique for the slender species comparison. DoS scores (for category all genes) are based on comparisons between the species *G. willdenowi* (stout) and *G. hofrichteri* (slender).

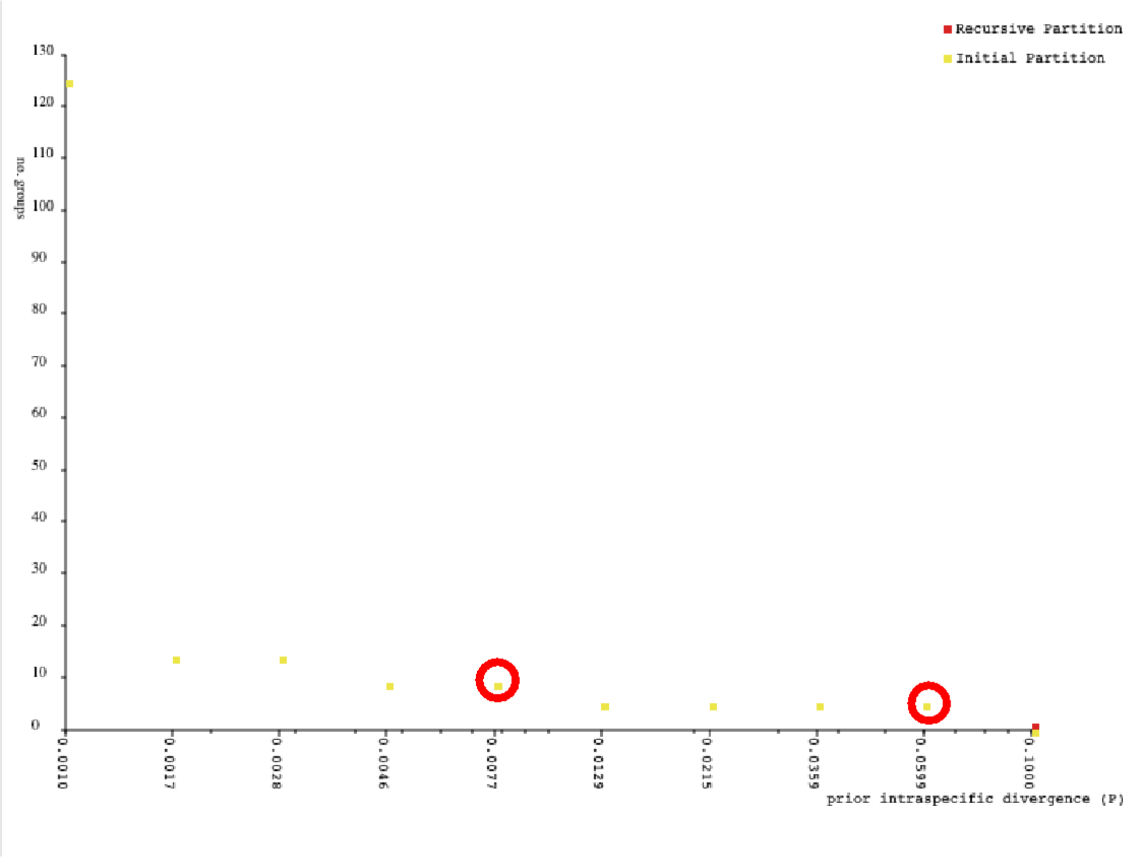


**Figure S19.**
The corresponding clusters for the two red encircled barcoding thresholds (corresponding to values 0.011 and 0.063) as obtained for Barcode gap distances inferred in ABGD using the K80 model.
